# Leukemia stemness and co-occurring mutations drive resistance to IDH inhibitors in acute myeloid leukemia

**DOI:** 10.1101/2020.10.27.357111

**Authors:** Feng Wang, Kiyomi Morita, Courtney D DiNardo, Ken Furudate, Tomoyuki Tanaka, Yuanqing Yan, Keyur P. Patel, Kyle J. MacBeth, Bin Wu, Guowen Liu, Mark Frattini, Jairo A. Matthews, Latasha D. Little, Curtis Gumbs, Xingzhi Song, Jianhua Zhang, Erika J. Thompson, Tapan M. Kadia, Guillermo Garcia-Manero, Elias Jabbour, Farhad Ravandi, Kapil Bhalla, Marina Konopleva, Hagop M. Kantarjian, P. Andrew Futreal, Koichi Takahashi

## Abstract

Allosteric inhibitors of mutant IDH1 or IDH2 induce terminal differentiation of the mutant leukemic blasts and provide durable clinical responses in approximately 40% of acute myeloid leukemia (AML) patients with the mutations. However, primary resistance and acquired resistance to the drugs are major clinical issues. To understand the molecular underpinnings of clinical resistance to IDH inhibitors (IDHi), we performed multipronged genomic analyses (DNA sequencing, RNA sequencing and cytosine methylation profiling) in longitudinally collected specimens from 68 IDH1- *or* IDH2-mutant AML patients treated with the inhibitors. The analysis revealed that leukemia stemness is a major driver of primary resistance to IDHi, whereas selection of mutations in *RUNX1/CEBPA* or *RAS-RTK* pathway genes was the main driver of acquired resistance to IDHi, along with *BCOR*, homologous *IDH* gene, and *TET2*. These data suggest that targeting stemness and certain high-risk co-occurring mutations may overcome resistance to IDHi in AML.

## Introduction

Somatic mutations in isocitrate dehydrogenase 1 and 2 (*IDH1 a*nd *IDH2*) can be detected in approximately 20% of patients with acute myeloid leukemia (AML) ^1^. Mutations are almost exclusively found in the Arg132 (R132) residue in IDH1 and Arg140 (R140) or Arg172 (R172) residues in IDH2. Wild-type IDH1 and IDH2 catalyze the oxidative decarboxylation of isocitrate to produce α-ketoglutarate (α-KG). On the other hand, mutant IDH1 and IDH2 acquire neomorphic catalytic activity and produce an oncometabolite, (R)-2-hydroxyglutarate [(R)-2HG or 2HG] ^2,3^, which competitively inhibits α-KG-dependent enzymes such as the ten-eleven translocation (TET) family of DNA hydroxylases, lysine histone demethylases, and prolyl hydroxylases ^4–6^. As a result, IDH-mutant AML exhibits CpG hypermethylated phenotype (CIMP) and increased histone methylation, leading to an aberrant gene expression profile and differentiation arrest ^7,8^.

Allosteric inhibitors to IDH mutant proteins (e.g. enasidenib for mutant IDH2 and ivosidenib for mutant IDH1) suppress 2HG production ^9^ and demonstrate an approximately 40% overall response rate in patients with IDH1- or IDH2- mutant relapsed and refractory AML ^10,11^. Clinical responders to the inhibitors show improvement in tri-lineage hematopoiesis and reduction of leukemic blasts. In the majority of the responders, *IDH* mutations are stably detected in matured neutrophils, indicating that the clinical response to the inhibitors is mediated by the terminal differentiation of leukemic blasts ^9^. This mechanism of action is consistent with the observations in preclinical models^12,13^ and patient-derived xenograft models^14^, as well as in longitudinally profiled hematopoietic stem cell populations from patients who responded to enasidenib ^15^. While the clinical response to IDHi can be durable, primary and secondary resistance to single-agent therapy are major clinical challenges ^10,11^. In a phase 2 study of enasidenib, co-occurrence of *NRAS* mutations or high co-mutation burden were associated with a poor response to the drug ^9^. Intlekofer and colleagues reported 3 cases that developed secondary resistance to enasidenib or ivosidenib ^16^. These cases acquired second-site mutations in the IDH2 dimer interface (p.Q316E and p.I391M) or IDH1 p.S280F, which were predicted to interfere with the IDHi binding. The same group of investigators also reported 4 cases of “IDH isoform switching”, which refers to the emergence of the mutation in homologous *IDH* gene counterpart during the inhibition of the other IDH mutant (e.g., emergence of *IDH1* mutation during IDH2 inhibition, and vice versa) ^17^. Additionally, Quek and colleagues studied paired samples at baseline and relapse in 11 AML patients treated with enasidenib ^15^. They did not find the second-site mutations but observed diverse patterns of clonal dynamics (including IDH isoform switching) or selection of sub-clones associated with the relapse.

While the data from the small case series are accumulating, the entire landscape of clonal heterogeneity and its association with IDHi resistance has not been elucidated. Moreover, the evidence accumulated so far has been restricted to the association between gene mutations and IDHi resistance. To what extent, DNA methylation changes or gene expression profiles are associated with clinical resistance to IDHi is not well understood.

Here, we performed an integrated genomic analysis combining DNA sequencing, RNA sequencing, and methylation profiling microarray on bone marrow samples collected longitudinally from AML patients treated with IDHi and described genetic and epigenetic correlates of response to IDHi. The analysis revealed that gene expression signatures with stemness is associated with primary resistance to IDHi, whereas selection of the resistant mutations plays role in acquired resistance to the drugs. These data add novel insights into the resistance mechanisms of IDHi in AML.

## Results

### Clinical characteristics of the studied patients

Clinical characteristics of the 68 patients are provided in the Figure 1A. Thirty-nine (57%) patients were *IDH2*-mutated, 28 (38%) were *IDH1*-mutated, and 1 (1%) had both mutations. Thirty-eight (56%) patients were treated with enasidenib, 22 (32%) with ivosidenib, 7 (10%) with IDH305, and 1 (1%) with AG-881. The selection of patients was solely based on availability of samples, resulting with a cohort of 38 clinical responders (56%) and 29 nonresponders (43%) (response not evaluable in 1 patient). Overall response rate (ORR) was not significantly different between patients treated with IDH1 inhibitors (ORR 66%) and IDH2 inhibitors (ORR 49%) (P = 0.204). Among the 38 responders, 25 patients relapsed after a median duration of response of 6.6 months (interquartile range: 3.6-13.5). Compared with the samples that were not analyzed in this study (due to the lack of sample availability), the studied cohort were older and contained more responders (Supplemental Table S1).

**Figure 1.**
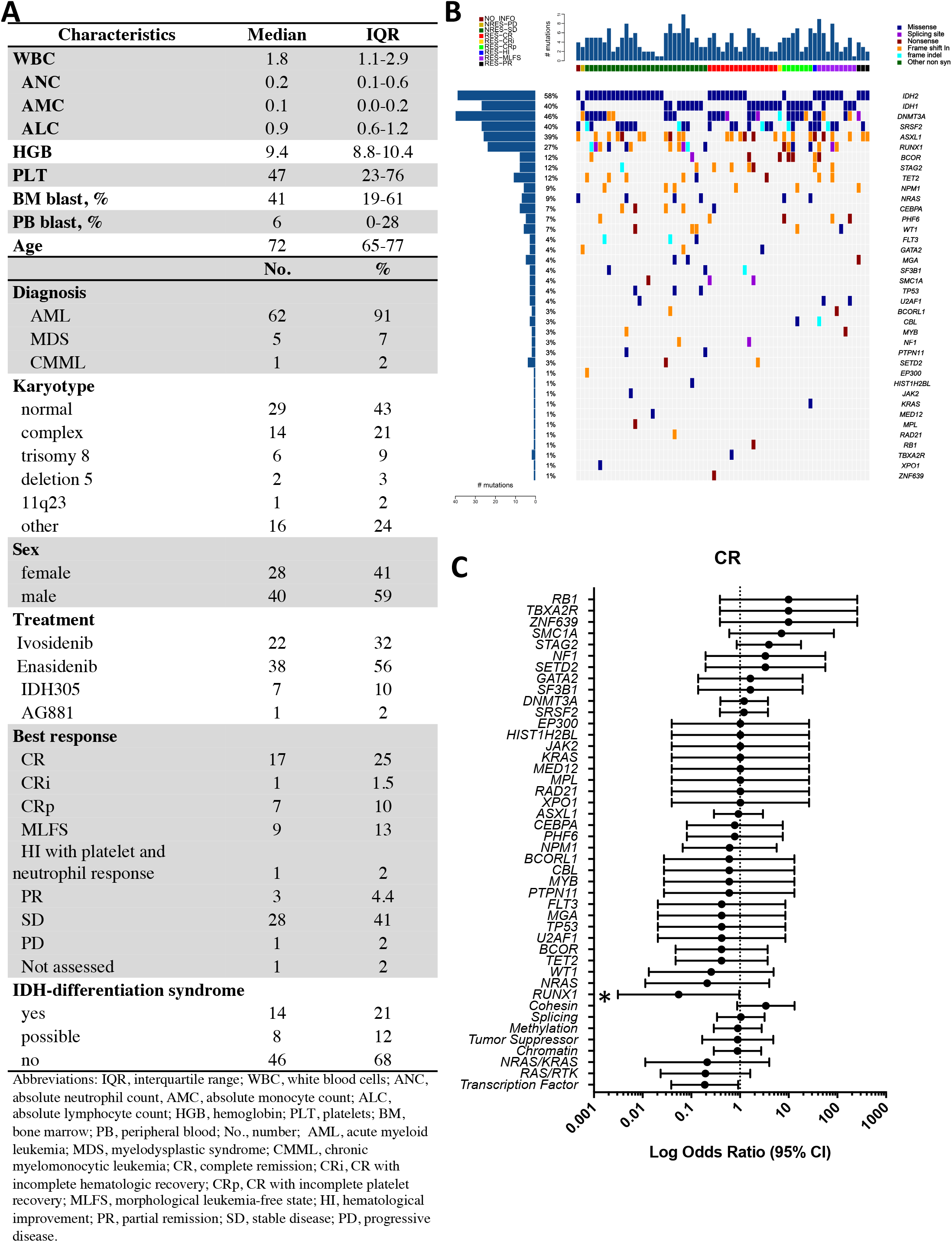
Clinical and mutational landscape of IDHi-treated AML patients and their association with clinical response. **(A)** Clinical characteristics of the 68 patients treated with IDH inhibitors. **(B)** Landscape of high-confidence somatic mutations detected in baseline samples by sequencing with a 295-gene panel. Legend for the best response is located at the top left while legend for the mutation classification is located at the top right. Baseline mutation data is available for 67 patients. **(C)** Forrest plot showing enrichment of the mutations at baseline against Complete Remission (CR) by logarithmic odds ratio. *P < 0.05. Circles represent odds ratios. The error bars represent 95% confidence interval of odds ratio. Baseline mutation data is available for 67 patients, out of which 16 achieved CR.

### Co-occurring RUNX1 or RAS signaling mutations are associated with primary resistance to IDH inhibitors

Targeted deep sequencing of pre-treatment samples identified 294 high-confidence somatic mutations (200 single-nucleotide variants [SNVs] and 94 small insertions and deletions [indels]) in 38 cancer genes (Figure 1B). Mutations that co-occurred with *IDH1/2* mutations were most frequently found in *DNMT3A* (N = 31, 46%), *SRSF2* (N = 27, 40%), *ASXL1* (N = 26, 39%), and *RUNX1* (N = 18, 27%). Relative timing of the mutation accrual was inferred based on the estimated cancer cell fraction (CCF) of the co-occurring mutations. In relative to *IDH1/2* mutations, mutations in *SRSF2, U2AF1, DNMT3A*, and *RUNX1* were predicted to have occurred earlier, whereas mutations in oncogenic *RAS* pathway genes (*NF1, PTPN11, CBL*, and *NRAS*) were likely acquired later (i.e., subclonal, Figure S1).

The analysis of co-occurring mutations and clinical response revealed that patients with concurrent *RUNX1* mutations had significantly inferior complete remission (CR) rate, and patients with concurrent *NRAS* mutations, previously associated with a poor CR rate with enasidenib, had a trend toward lower CR rate (Figure 1C and Figure S2). Of note, none of the patients with co-occurring *TP53* (N = 3) or *FLT3* mutations (N = 2) responded to the therapy while the association was not statistically significant due to the small number of cases. When genes were grouped with functional pathways, co-occurring mutations in hematopoietic differentiation transcription factor (TF) genes (*RUNX1, CEBPA*, and *GATA2*) were associated with a significantly worse CR rate and mutations in *RAS-RTK* pathway (*NRAS*, *KRAS*, *CBL*, *NF1, PTPN11*, and *FLT3*) had a trend toward worse CR rate (Figure 1C). In contrast, cooccurring mutations in cohesin genes (*STAG2, SMC1A*, and *RAD21*) were associated with a trend toward better response (Figure 1C). In the current cohort, we did not find a significant association between treatment response and the total number of co-occurring mutations (Figure S3).

### Leukemia stemness is associated with primary resistance to IDHi

Consensus *k*-means clustering of promoter methylation profiles in pre-treatment samples revealed two major clusters; Cluster 1 with relative hypomethylation and Cluster 2 with relative hypermethylation (Figure 2A-2B and Figure S4). *DNMT3A* mutations were significantly more frequent in Cluster 1 compared to Cluster 2 (Figure 2C), which likely accounts for the relative hypomethylation of the cluster, since co-occurrence of *DNMT3A* mutations with *IDH* mutations has been shown to cause methylation antagonism^18^. None of the other driver mutations had a significant correlation with the methylation-based clusters (Figure 2C).

**Figure 2.**
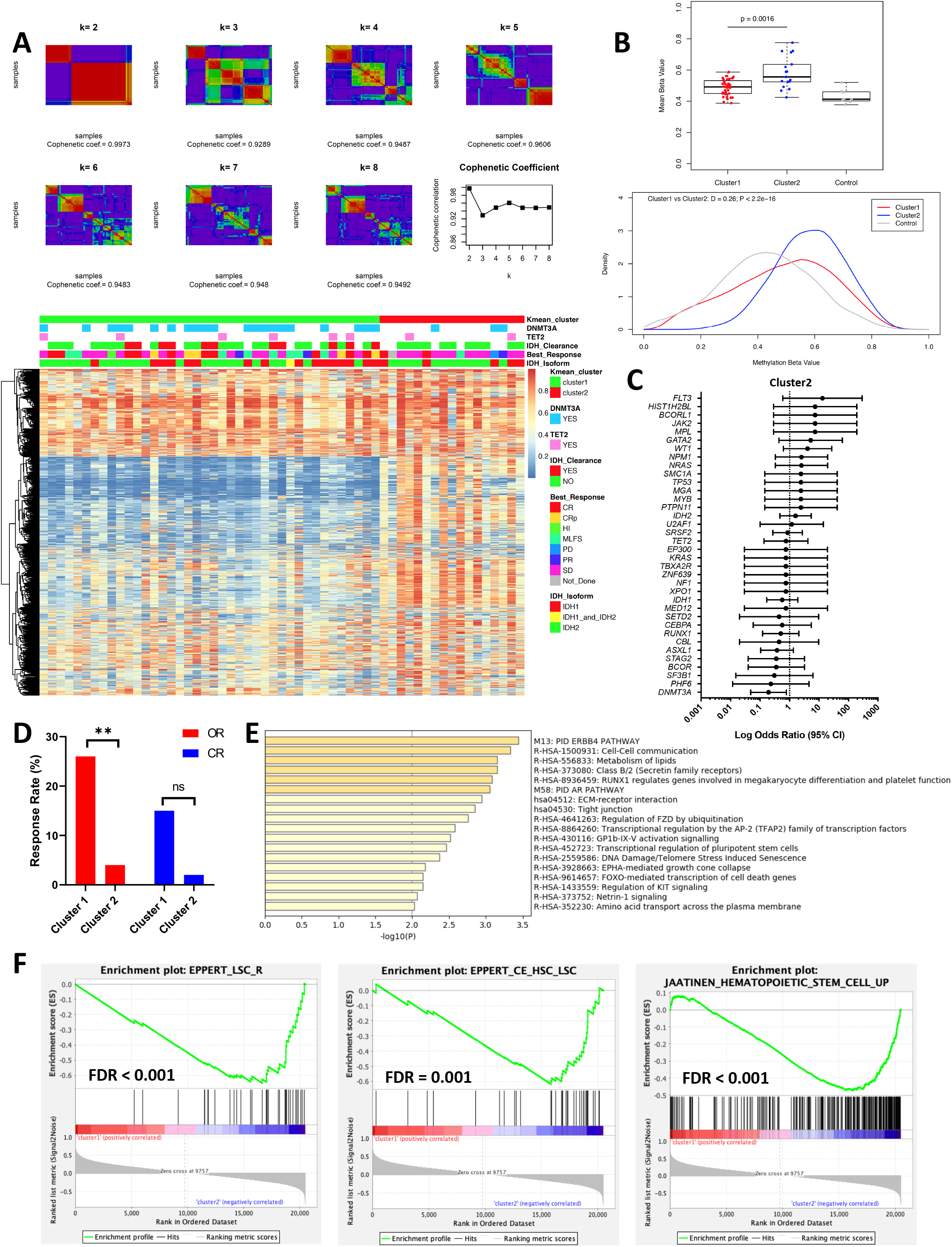
Analysis of DNA methylation at baseline samples reveals two distinct clusters associated with treatment response. **(A)** Consensus k-mean clustering of promoter methylation data at baseline revealed two distinct clusters. Methylation data is based on methylation beta value. Promoter CpG probes from top 1% most variably methylated CpG probes were selected for the analysis. Responders were defined as patients achieved best response of CR, CRp, MLFS, PR and HI. Non-responders were defined as patients achieved best response of PD and SD. **(B)** Top box plot comparing mean methylation beta value of top 1% most variably methylated CpGs among Cluster 1 baseline (N=40) and Cluster 2 baseline (N=17) samples. IDH1/2 wild type AML samples (N=8) are used as control. Bottom density distribution of top 1% most variably methylated CpG probes with methylation beta values comparing Cluster 1 baseline and Cluster 2 samples. Kolmogorov–Smirnov test D and P values are shown. IDH1/2 wild type AML samples (N=8) are used as control. **(C)** Forrest plot showing enrichment of the mutations at baseline against being in Cluster 2 by logarithmic odds ratio. *P < 0.05. Circles represent odds ratios. The error bars represent 95% confidence interval of odds ratio. Baseline mutation data with clustering information is available for 57 patients, out of which 17 is in Cluster 2. **(D)** Bar plot comparing the Overall Response (OR) and CR rate between Cluster 1 and Cluster 2 patients. ** P<0.01 **(E)** Metascape analysis of hypermethylated promoter DMPs **(F)** Gene Set Enrichment Analysis (GSEA) comparing gene expression profiles between the two clusters revealed upregulation of genes associated with leukemia stem cells (LSCs) in Cluster 2.

Notably, Cluster 2 (hypermethylated cluster) was associated with significantly poor response to IDHi (Figure 2D). The analysis of differentially methylated probes (DMP) between the two clusters showed that promoters in genes related to hematopoietic differentiation, such as RUNX1 targets, transcriptional regulation of stem cells, and KIT signaling, are significantly hypermethylated in Cluster 2 (Figure 2E). While the expression of more than half of the DMP genes were downregulated in Cluster 2 compared to Cluster 1, there was no consistent trend for promoter hypermethylation and gene downregulation in DMP genes (Figure S5).

We then analyzed the difference in gene expression profiles between the two clusters. Gene Set Enrichment Analysis (GSEA) comparing gene expression profiles between the two clusters revealed upregulation of genes associated with leukemia stem cells (LSCs) in Cluster 2 (Figure 2F). To further explore the molecular drivers of Cluster 2 phenotype, we performed NetBID analysis, a data-driven network-based Bayesian inference that identifies hidden drivers in a given transcriptome^19^. Among the top driver transcription factors genes enriched in Cluster 2 included *FOXC1*, which is one of the critical regulators of LSC function (Figure 3A)^20^. Additional drivers identified for Cluster 2 included *CD99* and *CDK6*, both of which encode essential signaling proteins for LSC (Figure 3B)^21,22^. In addition, *DNMT3A* was identified as one of the drivers in Cluster 2, that is consistent with the lack of *DNMT3A* mutations in the cluster because *DNMT3A* mutations are generally loss of function mutations (Figure 3B). Together, these data suggest that Cluster 2 is enriched with samples manifesting increased stemness, which might be associated with resistance to IDHi.

**Figure 3.**
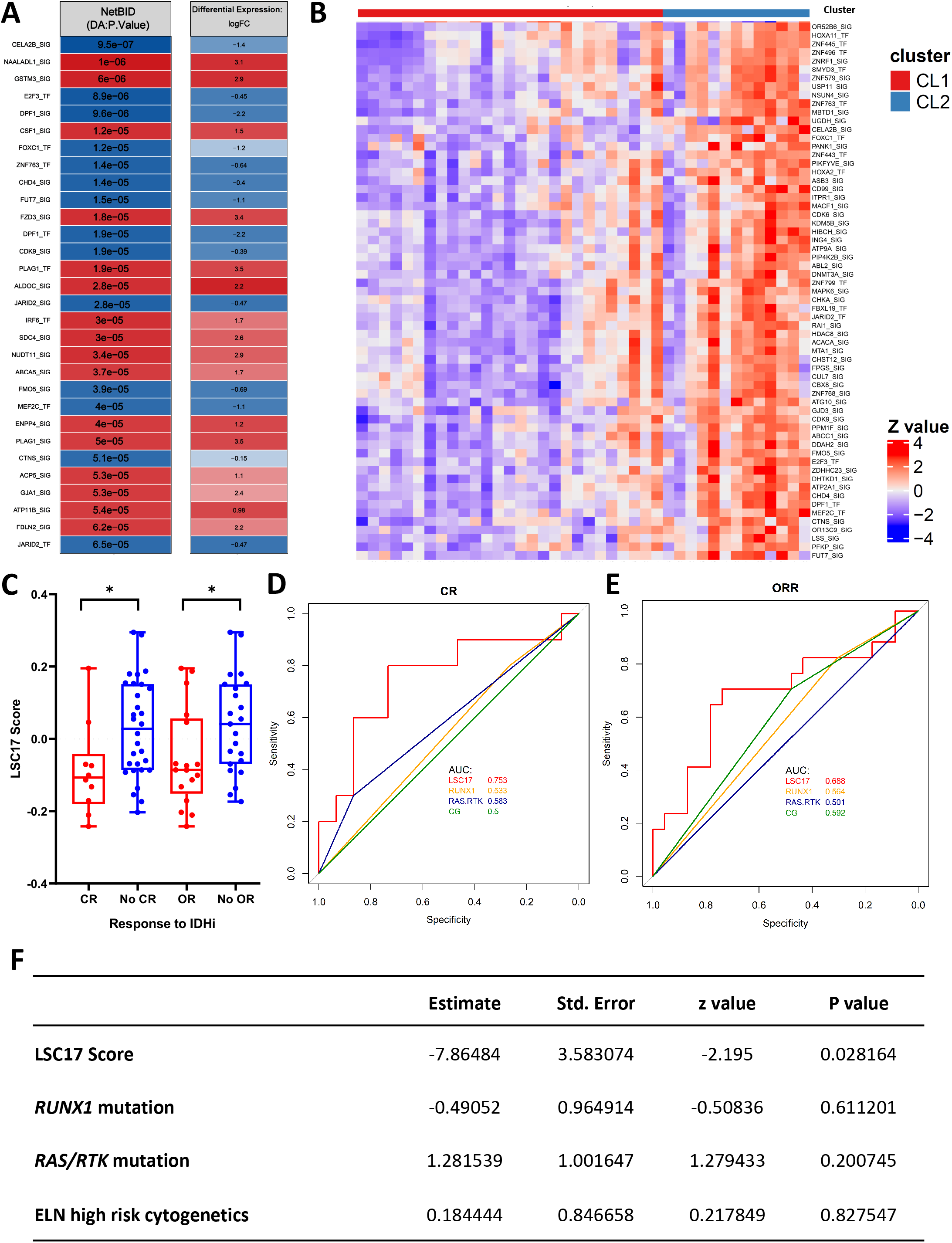
Leukemia stemness is associated with primary resistance to IDHi. **(A)** List of top driver transcription factors (TF) and signaling genes (SIG) identified by NetBID2 analysis by comparing gene expression profiles between Cluster 1 and Cluster 2 (P < 0.01 was used as the significance cutoff). Drivers identified for Cluster 1 and Cluster 2 are colored with red and blue, respectively. (**B**) Heatmap of NetBID-based activity of top drivers in Cluster 2. Samples in Cluster 1 (CL1) are labeled as red while Cluster 2 (CL2) are labeled as blue. (**C**) LSC17 score was calculated for each baseline sample and compared between patients achieving CR vs. not and OR vs. not. * P<0.05. (**D-E**) Receiver operating curve (ROC) for predicting CR or OR with LSC17 score, *RUNX1* mutation status, *RAS-RTK* mutation status, and ELN cytogenetic risk classification. **(F)** Multi-logistic regression analysis against CR by considering following variables: LSC17 score (as a continuous variable), *RUNX1* mutation status (mutated vs. wild type), *RAS-RTK* mutation status (mutated vs. wild type), and ELN high rick cytogenetics classification (high risk vs. others).

To determine the association between leukemia stemness and IDHi resistance, we calculated the 17-gene LSC score (LSC17) for each sample, which has been associated with leukemia stemness and chemoresistance in AML^23^. Non-responders to IDHi had a significantly higher LSC17 compared with responders (Figure 3C). LSC17 predicted response to IDHi (CR) with AUROC (area under the curve of receiver operating curve) of 0.75 (P = 0.018), which was better than the predictability of ELN cytogenetic risks (AUROC = 0.5), *RUNX1* mutations (AUROC = 0.54), or *RAS-RTK* mutations (AUROC = 0.58) (Figure 3D-E). Multi-logistic regression analysis also showed that LSC17 was the significant covariate predicting response to IDHi (Figure 3F). Collectively, these data indicate increased stemness as one of the mechanisms of primary resistance and the stemness score as a potential predictive biomarker for IDHi response.

### DNA methylation changes after IDHi

We then analyzed the changes in DNA methylation after IDHi therapy. While there were some heterogeneities among samples, overall, we observed significant demethylation after IDHi (Figure 4A). Demethylation was observed in samples regardless of the methylation-based clusters (Cluster 1 vs. 2) or treatment response. Consistent with this, plasma 2HG was also suppressed after IDHi in most of the patients regardless of the clusters and treatment responses (Figure 4B). While Cluster 1 and Cluster 2 both exhibited incremental demethylation after the therapy, Cluster 2 remained relatively hypermethylated after the therapy compared to Cluster 1 (Figure 4A). The analysis of methylation changes in individual CpGs revealed that the same set of CpGs were demethylated between Cluster 1 and Cluster 2 (Figure 4C). Consistent with this, post-treatment DMPs between the two clusters were largely the same with those at baseline, with most of the DMPs remained hypermethylated in Cluster 2 (Figure 4D-E). The same trend was observed when we compared the methylation changes in individual CpGs between responders and non-responders (Figure 4F). GSEA comparing gene expression profiles of post-treatment samples in Cluster 1 and 2 showed that LSC-associated genes are still upregulated in Cluster 2 (Figure 4G), suggesting that the stemness is not reversed by IDHi. Collectively, these results suggest that incremental changes in DNA methylation are likely the consequence of 2HG suppression by IDHi therapy and does not necessarily contribute to the clinical response.

**Figure 4.**
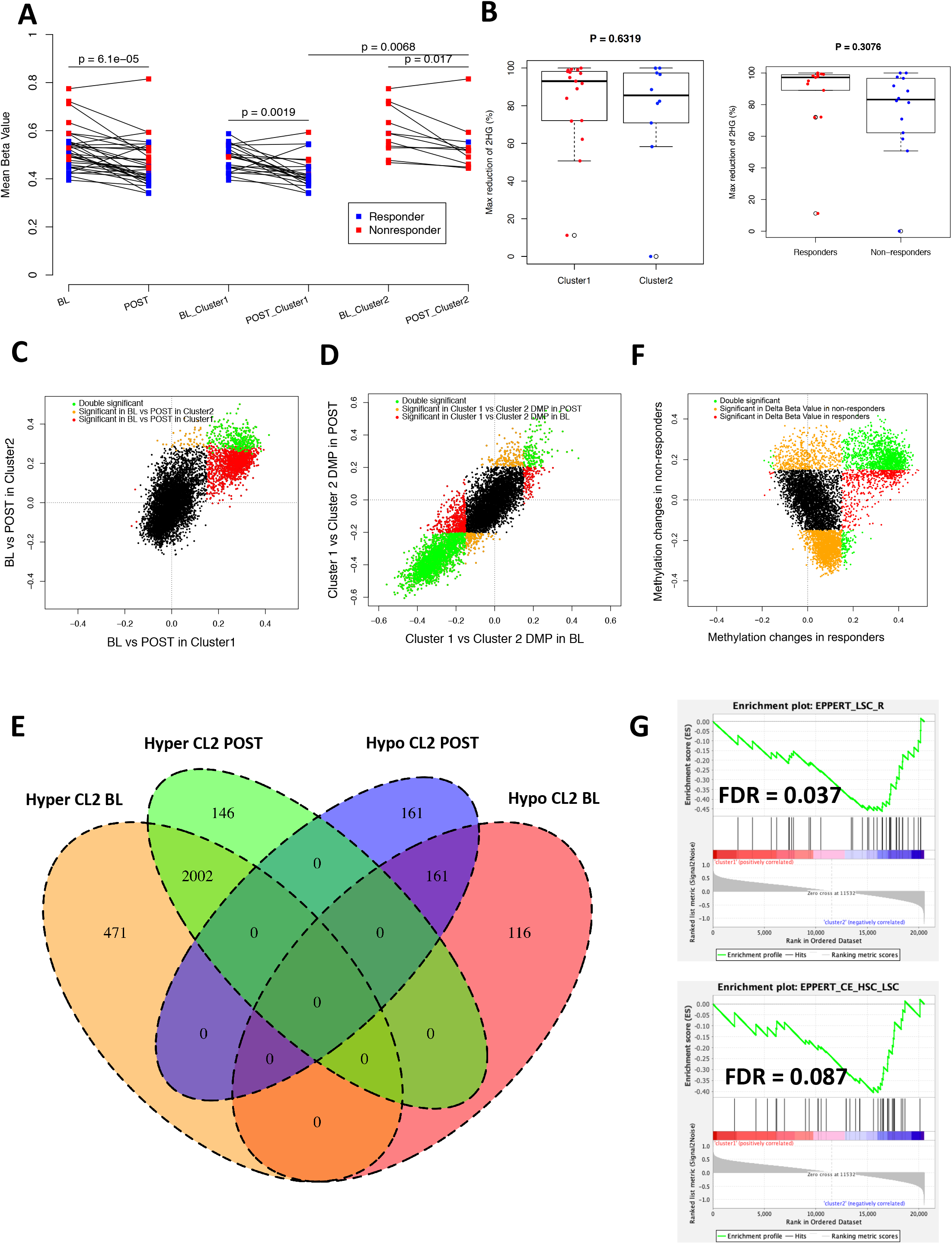
DNA methylation changes after IDHi. **(A)** Longitudinal trend of methylation level at Baseline (BL) and Post-treatment (POST) for all, Cluster 1 and Cluster 2 patients. Responders and non-responders are color coded. **(B)** Box plot showing maximum reduction of plasma 2HG levels after IDHi treatment (%) in Cluster 1 and Cluster 2 patients as well as responders and nonresponders. **(C)** Scatterplot showing correlation of the longitudinal methylation changes between Cluster 1 and Cluster 2 patients. Each dot represents a CpG probe and was colored based on its significance in the longitudinal differentially methylation test in either Cluster 1 or Cluster 2 patients. The X axis represents the differential methylation level between BL and POST samples (i.e., beta value at BL minus beta value at POST) in Cluster 1 patients and the Y axis represents the differential methylation level between BL and POST samples in Cluster 2 patients. **(D)** Scatterplot showing correlation of the inter-cluster methylation differences between BL and POST time points. Each dot represents a CpG probe and was colored based on its significance in the inter-cluster differentially methylation test at either BL or POST time points. The X axis represents the differential methylation level between Cluster 1 and Cluster 2 in BL samples (i.e., beta value of Cluster 1 minus Cluster 2) and the Y axis represents the differential methylation level between Cluster 1 and Cluster 2 in POST samples. (**E**) Venn diagram showing the overlapped DMPs between Cluster 1 and Cluster 2 at baseline (BL) and post-treatment (POST). Among 2619 DMPs hypermethylated in Cluster 2, 2002 overlapped between BL and POST, suggesting that most of the hypermethylated DMPs in Cluster 2 were the same before and after treatment. (**F**) Scatterplot showing correlation of the longitudinal methylation changes between responders and non-responders. Each dot represents a CpG probe and was colored based on its significance in the longitudinal differentially methylation test in either responders or nonresponders. The X axis represents the differential methylation level between baseline and response samples in responders (i.e., beta value at BL minus beta value at POST) and the Y axis represents the differential methylation level between baseline and non-response samples in nonresponders. (**G**) GSEA analysis comparing gene expression of post-treatment samples between Cluster 1 and Cluster 2 showed that LSC genes are still upregulated in Cluster 2 post-treatment.

### Clonal selection of driver mutations frequently accompanies relapse after IDHi

We then investigated the mechanisms of acquired resistance to IDHi by analyzing mutational changes in longitudinal samples collected after IDHi therapy. With regards to the *IDH* mutations, variant allele frequency (VAF) of the mutations stayed unchanged in 73% of the responders (N=22), whereas 27% of the responders (N=8) had a substantial reduction (>= 75% decrease of VAF) or clearance of the mutations at response (Figure S6). The baseline VAF or the types of *IDH* mutations did not predict the clearance of the mutations (Figure S7). Also, there was no correlation between *IDH* mutation clearance and the patterns of co-occurring mutations (Figure S8). We did not observe a significant difference in survival between patients who cleared the *IDH* mutation and who did not (Figure S9). In non-responders, *IDH* VAF were mostly unchanged but 4 patients had a substantial reduction on therapy (Figure S10).

Co-occurring mutations demonstrated variable dynamics during therapy. Emergence of previously undetectable mutations or selection of subclonal mutations frequently accompanied the relapse or disease progression (Figure 5A). Among the 23 patients with pre-treatment and relapse pairs, emerging or selected mutations were detected in 19 (83%) patients at the time of relapse (Figure 5A). Mutations that were frequently acquired or selected at relapse involved *RUNX1* and *BCOR* in 5 cases each, followed by *KRAS* and *NRAS* in 3 cases each (Figure 5B). IDH dimer-interface mutations were not detected in this cohort. Isoform switching occurred in 1 case. Overall, relapse-associated mutations involved *RAS-RTK* pathway (39%), chromatin structure (35%), hematopoietic transcription factors (30%), and DNA methylation pathways (22%) (Figure 5B). These relapse-associated mutations were remarkably similar to those associated with poor initial response to IDHi (Figure 1B and Figure 1C), highlighting their crucial role in the clinical resistance of IDHi. To determine whether the relapse-associated mutations are part of or independent of *IDH* mutations, we performed a single-cell DNA sequencing (scDNA-seq) in a subset of relapsed samples. In UPN2394529, the relapse was associated with emerging *KRAS* p.Q61H and *NRAS* p.G12S mutations (Figure 5C). The scDNA-seq revealed that the emerging *NRAS* and *KRAS* mutations were independent of *IDH2* mutation, indicating that non IDH-mutant clones were driving the relapse in this case (Figure 5E). In contrast, in UPN2297707, the relapse was accompanied with the selection of *RUNX1* p.K152fs mutation, which co-occurred with the *IDH1* mutation (Figure 5D and Figure 5F).

**Figure 5.**
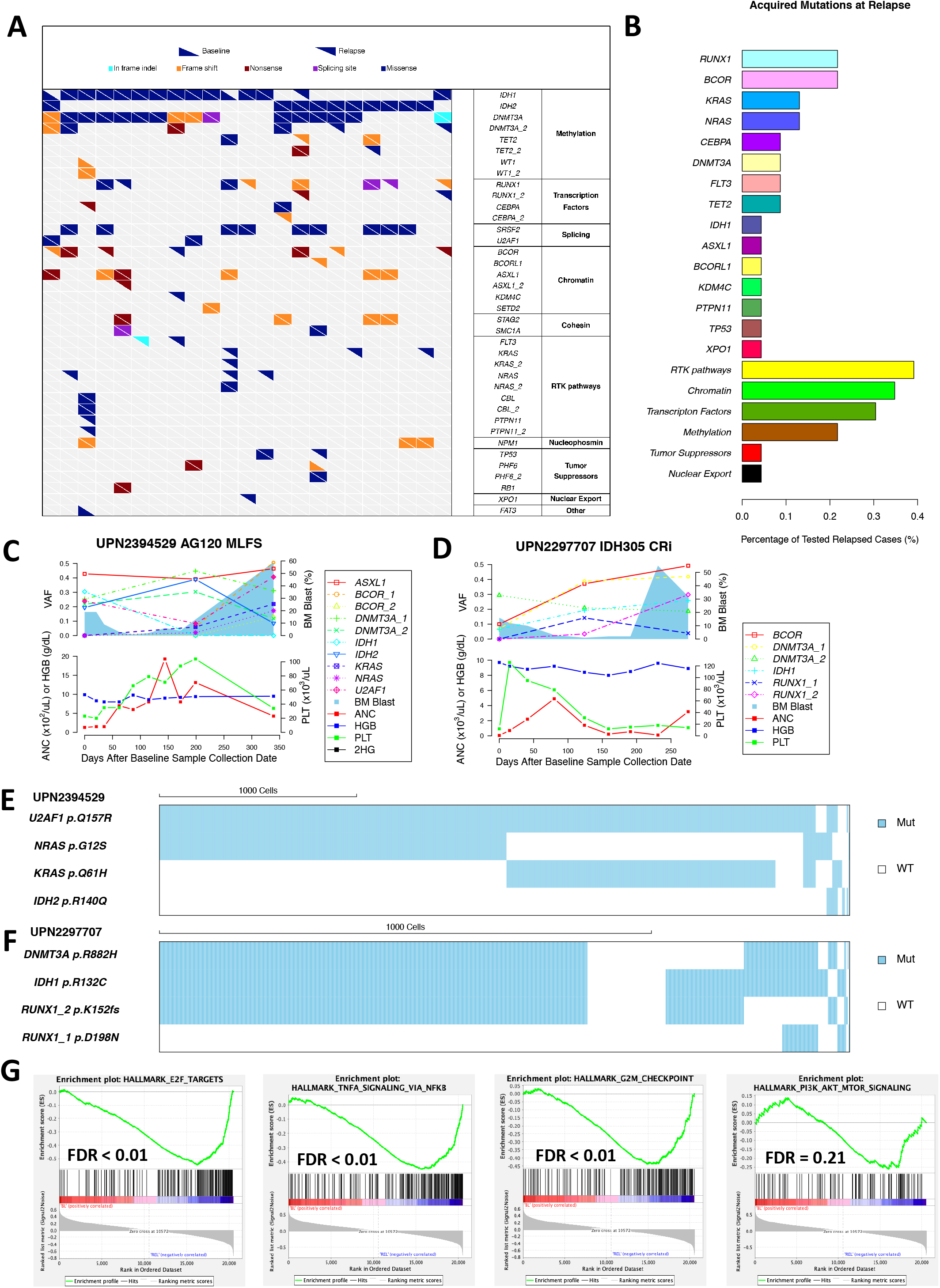
Selection of resistant mutations accompanies relapse after IDHi. **(A)** Longitudinal mutation landscape plot showing mutation acquisitions in 19 out of the 23 tested relapsed cases. Each column represents an individual case with differentially shaped triangles representing mutation status in either baseline or relapse. **(B)** Bar plot showing percentage of tested relapsed cases with acquired mutations in various genes and pathways. **(C-D)** The longitudinal trajectory of mutation VAFs, bone marrow (BM) blast counts, absolute neutrophil count (ANC), Hemoglobin (HGB) counts and Platelet (PLT) counts in UPN2394529 (C) and UPN2297707 (D). Line plots show mutation VAFs and ANC/HGB/PLT counts. Blue shades represent BM blast counts. **(E-F)** Single cell landscape of selected mutations in UPN2394529 (E) and UPN2297707 (F). Each column represents one individual cell. 1000 cells scale bar is shown on the top left. (**G**) GSEA comparing gene expression data from RNA sequencing between baseline and relapse samples showing significant enrichment of E2F targets, TNF alpha signaling via NF-kappa B, G2M checkpoint, and PI3K, AKT, MTOR signaling pathways in relapse samples.

GSEA comparing the gene expression profiles between pre-treatment and relapse pairs showed the enrichment of genes downregulated in LSC in relapsed samples (Figure S11), which contrasts to the samples with primary resistance, indicating for a difference between primary resistance and acquired resistance. Instead, relapsed samples were associated with upregulation of genes in E2F targets, TNF alpha signaling via NF-kappa B, G2M checkpoint, and PI3K, AKT, MTOR signaling pathway genes, that are consistent with the frequent acquisition of *RAS-RTK* pathway mutations at relapse (Figure 5G). Collectively, these results underscore the role of cooccurring mutations, particularly *RUNX1* and *RAS-RTK* pathway mutations in acquired resistance to IDHi, and that the co-occurring mutations can be part of or independent of IDH-mutant clones.

### Mapping genetic and epigenetic evolution during IDHi therapy in individual cases

The heterogeneity in genetic evolution and methylation changes after IDHi prompted us to investigate the dynamic changes in genome and epigenome during IDHi therapy at individual patient-level. Sixteen patients (5 responders and 11 non-responders) had a set of multidimensional data available at longitudinal timepoints to map the evolution of somatic mutations and DNA methylation along with clinical parameters and plasma 2HG in individual cases. This analysis identified three major patterns of epigenetic evolution in responders, which correlated with the underlying genetic evolution.

In the first pattern, IDHi effectively suppressed plasma 2HG and bone marrow DNA methylation level at response and the suppression of both markers continued at relapse. The pattern was observed in 2 caess, UPN1825001 and UPN2463247, of which the relapse was associated with growing *KRAS* mutations. In both cases, plasma 2HG remained suppressed at the time of disease progression, which correlated with sustained suppression of DNA methylation (Figure 6A-6B). In the second pattern, IDHi similarly suppressed both plasma 2HG and bone marrow DNA methylation at response, however, at relapse, we observed de-suppression of DNA methylation while plasma 2HG remained low. This pattern was associated with emerging *TET2* mutations at relapse, which is consistent with the role of *TET2* mutation in causing hypermethylation phenotype (observed in UPN2297625 and UPN2620771; Figure 6C-6D). In both pattern 1 and 2, IDHi remained functionally active at relapse (i.e., ongoing 2HG suppression).

**Figure 6.**
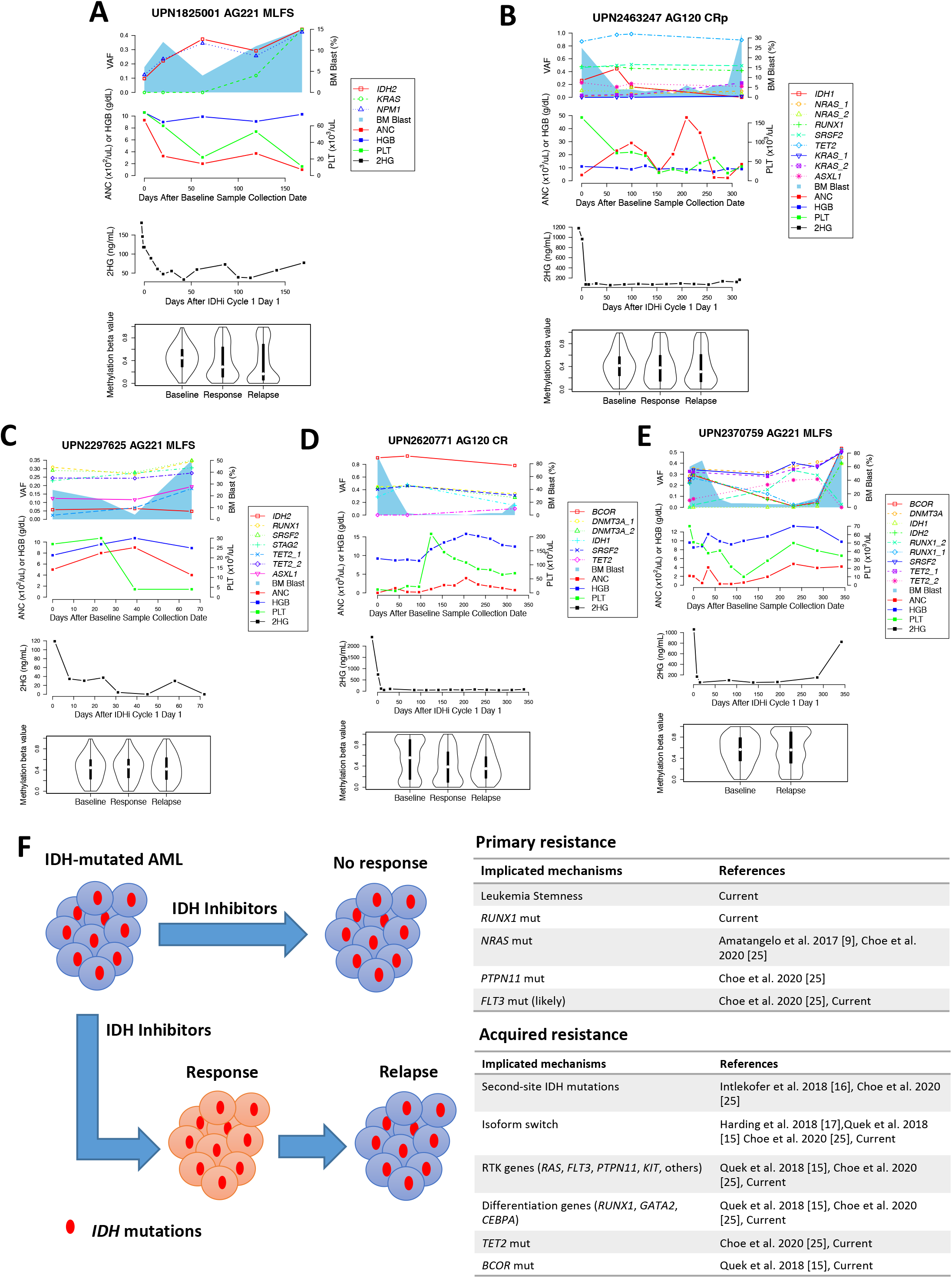
Heterogeneous patterns of genetic and epigenetic evolution in AML patients treated with IDHi. **(A-E)** Multi-dimensional longitudinal plot of mutation VAFs, bone marrow (BM) blast counts, absolute neutrophil count (ANC), Hemoglobin (HGB) counts, Platelet (PLT) counts, 2HG level and DNA methylation level in UPN1825001 (A), UPN2463247 (B), UPN2297625 (C), UPN2620771 (D), and UPN2370759 (E). Line plots show mutation VAFs, ANC/HGB/PLT counts and 2HG level. Blue shades represent BM blast counts. Violin plots show the methylation distribution. (**F**) A summary of available evidence for the association between molecular alterations and IDHi resistance including the result from the current study.

The third pattern was observed in one case UPN2370759, which was consistent with “IDH isoform switching” previously described ^17^. The case initially had *IDH2* mutation and was treated with AG221, however, the relapse was associated with an emergence of *IDH1* p.R132C mutation. In this case, both plasma 2HG and methylation levels increased at relapse, which is consistent with the emergence of *IDH1* mutation during IDH2 inhibition (Figure 6E). The *IDH1* p.R132C mutation that emerged was not detectable at baseline by both targeted sequencing and the digital droplet PCR (ddPCR) assay (sensitivity 0.01%), making it more likely that the mutation was acquired *de novo* at relapse (Figure S12).

In non-responders, 9 of 11 samples showed co-suppression of plasma 2HG and DNA methylation after IDHi therapy while it did not lead to clinical response in these patients. In 2 cases, despite suppression of plasma 2HG, we did not observe demethylation. Underlying mechanisms of this discrepancy is not clear. Genetic and epigenetic evolution of non-responders are shown Figure S13.

## Discussion

Using a multipronged genomic analysis on longitudinally collected samples from the clinical trials, we studied genetic and epigenetic correlates of response to IDHi in AML. While confirming previous findings about the role of certain co-occurring mutations (*RAS* and *RUNX1*) in primary resistance to IDHi^9^, we additionally and notably revealed that leukemia stemness is associated with IDHi primary resistance. In the current cohort, higher LSC17 score was the strongest predictor of response to IDHi. Since the clinical activity of IDHi is driven by the induction of terminal differentiation of leukemic blast^14^, it is plausible that stemness phenotype causes inherent resistance to the differentiating mechanism of action of IDHi monotherapies. The underlying mechanisms driving stemness in IDH-mutant AML is unclear. Increased stemness was associated with hypermethylated phenotype (Cluster 2) in this cohort, which had a further association with the absence of co-occurring *DNMT3A* mutations. Whether the hypermethylation (or the lack of co-occurring *DNMT3A* mutations) is directly causing an increased stemness is not clear. While hypermethylated DMPs were enriched in promoters associated with hematopoietic differentiation genes, the effect on the corresponding gene expression was modest. Since *IDH*-mutations broadly affects methylation status including enhancers and histones, further investigation is needed to understand the mechanisms driving the stemness in *IDH*-mutated AML and the connection between the stemness and hypermethylation status. Nonetheless, our findings, while require validation in independent cohort, offer a possibility that stemness signatures may function as a predictive biomarker for IDHi response.

Co-occurring mutations and the selection of resistant mutations were also critical factors for IDHi resistance, particularly in the setting of acquired resistance^24^. There was no single dominant gene mutation associated with the resistance, however, the mutations implicated for the resistance were remarkably consistent in both primary and acquired resistance settings, underscoring their role in clinical resistance to IDHi. One of the major pathways affected by the mutations were hematopoietic differentiation transcription factors, particularly involving *RUNX1. RUNX1 co*-mutation(s) at baseline was associated with lower CR rate in our cohort. *RUNX1* mutations were also among the most frequently acquired or selected mutations at relapse. In addition, 4 of 5 patients with co-occurring *CEBPA* mutations did not respond to IDHi and the mutation were also acquired at relapse in 2 patients (Figure 5A). Since *RUNX1* and *CEBPA* both encode essential transcription factors for hematopoietic and myeloid differentiation, mutations in these genes likely abrogates differentiation signals induced by IDHi, thus contributing to the clinical resistance.

Mutations in the *RAS-RTK* pathway represent another major mechanism of the resistance. The association between co-occurring *RAS* pathway mutations and primary resistance to enasidenib or ivosidenib has been previously reported^9,25^. In our cohort, co-occurrence of *NRAS* mutations at baseline trended with a poor response to the IDHi therapy. Additionally, *NRAS* or *KRAS* mutations had been acquired at relapse in nearly 30% of the cases. Intriguingly, in at least one case that had acquired *KRAS* and *NRAS* mutations at relapse (UPN2394529), the mutations did not co-occur with *IDH* mutation by the single-cell sequencing, suggesting that selection of non-IDH clone can also drive relapse.

While co-occurring mutations in *RUNX1/CEBPA* or *RAS-RTK* genes were the major pathways to IDHi resistance, we also observed other less frequent but intriguing mechanisms. One was an acquired mutation in the homologous gene. The same phenomenon (described as “isoform switching”) was previously reported in 2 cases of AML treated with ivosidenib ^17^. This pattern was associated with an increase in plasma 2HG and DNA hyper-methylation at relapse. We also observed acquisition of *TET2* mutation as a likely IDHi resistance mechanism. In contrast with the isoform switching, these *TET2*-acquired cases showed continued suppression of plasma 2HG at relapse while DNA hypo-methylation did not occur. We also observed frequent acquisition of loss-of-function mutations in the *BCOR* gene at relapse. *BCOR* is part of non-canonical PRC1.1 complex, which acts as a transcription corepressor ^26^. It is not yet clear how loss of BCOR function contributes to IDHi resistance, but the data offer hypothesis that BCOR target genes may be involved in IDHi resistance. In our cohort, we did not observe the acquisition of second-site mutations at the dimer interface of IDH1/2 ^16^. This was also not found in 11 cases of post enasidenib relapse analyzed by Quek et al ^15^. A recent study by Choe et al. identified the second-site mutations in 14% of IDH1-mutant AML patients who relapsed after ivosidenib^25^. Although the dimer interface mutations in *IDH1/2* represent a compelling mechanism of acquired resistance to IDHi, further studies are needed to understand the true prevalence of this mechanism. We have reviewed and summarized the available evidence related to the molecular mechanisms of IDHi resistance in Figure 6F.

This study also analyzed dynamic changes in CpG methylation during IDHi therapy. The drug induced hypomethylation in bone marrow samples that is consistent with the suppression of 2HG (likely through the restoration of TET family protein activity). However, the incremental changes in DNA methylation occurred in the same CpGs among responders and non-responders, and there was no concordance between DNA methylation changes and clinical response. These data suggest that incremental changes in DNA methylation mirror the 2HG dynamics except in rare cases with co-occurring *TET2* mutations, and do not correlate with clinical response to IDHi. There are several limitations in our study. First, we could not independently validate the association between stemness signature and IDHi response. This finding needs to be confirmed in an independent cohort of AML patients treated with IDHi. Second, the sample size of this study was underpowered to capture rare molecular predictors of IDHi response, for example *FLT3* and *TP53*. Results from the several independent studies correlating gene mutations and IDHi response are now available, and meta-analysis of the combined dataset might reveal the entire landscape of gene mutations and their impact on IDHi response. Third, due to the limited amount of the available specimens, multi-omics analyses were not possible in all sample, leading to inconsistencies in data generation among samples (Figure S14). Fourth, our cohort included heterogeneous patient populations who were at the different stages of their disease and also included small number of patients with MDS/CMML. Also, different types of IDH inhibitors were given to the patients (enasidenib, ivosidenib, IDH305, and AG881). While we do not believe these heterogeneities affect overall conclusion of our study, our findings need to be validated in patients with more uniform characteristics. Lastly, the targeted DNA sequencing might have missed low VAF mutations for mutation clearance and clonal dynamics analysis. With all these limitations in mind, we believe that our study adds novel insights into genetic and epigenetic mechanisms of resistance to IDHi.

In summary, the molecular profiling of IDHi-treated AML samples revealed that leukemia stemness plays major role in primary resistance to the drug, whereas co-occurring mutations, particularly in hematopoietic transcription factor genes (*RUNX1* and *CEBPA*) and *RAS-RTK* genes, are critical factors for acquired resistance. These results suggest that novel strategies targeting stemness and co-occurring mutations may improve the therapeutic efficacy of IDHi in AML. The results from ongoing combination therapy trials (IDHi with azacitidine, cytarabine+daunorubicin, MEK inhibitor, or venetoclax) are warranted to understand how these approaches can overcome these resistance mechanisms.

## Methods

### Patients and samples

We studied 68 patients with relapsed or refractory myeloid malignancies (AML N =62, MDS N = 5, and CMML N =1) who received IDH inhibitor therapy in one of the 4 clinical trials conducted in our institution: NCT01915498 (enasidenib for *IDH2* mutated patients), NCT02074839 (ivosidenib for *IDH1* mutated patients), NCT02381886 (IDH305 for *IDH1* mutated patients), and NCT02481154 (AG-881 for *IDH1* or *IDH2* mutated patients). Selection of the studied patients was based on the sample availability alone. Bone marrow mononuclear cells were collected longitudinally (pre-treatment, post-treatment, and relapse) from the trial participants and were subject for the analyses. Clinical response to the therapy was determined by the clinical investigators and followed the modified 2003 International Working Group criteria^27^. We defined “responders” as having overall response to the therapy, which included CR, CR with incomplete hematologic or platelet recovery, partial response, and MLFS. In MDS patients, hematologic improvement was also considered as response. “Non-responders” were defined as having stable disease or progressive disease. Written informed consent for sample collection and analysis was obtained from all patients. The study protocols adhered to the Declaration of Helsinki and were approved by the Institutional Review Board at The University of Texas MD Anderson Cancer Center. Detailed information about the sample availability is shown in Figure S14.

### Targeted deep sequencing and data analysis

We used a SureSelect custom panel of 295 genes (Agilent Technologies, Santa Clara, CA) which are recurrently mutated in hematologic malignancies (Table S2). Details of the sequencing methods have been described previously ^28^. The same panel was used for all baseline and longitudinal samples. Briefly, genomic DNA was extracted from bone marrow mononuclear cells using an Autopure extractor (QIAGEN/Gentra, Valencia, CA). All longitudinal samples were analyzed by the same targeted panel sequencing. DNAs were fragmented and bait-captured in solution according to manufacturer’s protocols. Captured DNA libraries were then sequenced using a HiSeq 2000 sequencer (Illumina, San Diego, CA) with 76 base pair paired-end reads. The median of median depth of the targeted regions was 393x (IQR: 332-485x). Bioinformatic pipelines calling high-confidence somatic single nucleotide variants (SNVs) and indels from targeted capture DNA sequencing were described previously ^28^. Detailed description is available in supplementary material.

### Methylation array profiling and data analysis

DNA methylation analysis was performed using Illumina’s Infinium MethylationEPIC assay (EPIC) according to the manufacturer’s protocol as previously described ^29^. Data analysis was conducted using the ChAMP algorithm^30^ as previously described using default parameters. Briefly, The IDAT files were taken as input files and raw beta values were generated. Following initial quality check and probe filtering including removing all SNP-related probes ^31^ and all probes located in chromosome X and Y, the data were normalized using the BMIQ method^32^. Differential methylation analysis was performed by using the limma algorithm^33^.

### RNA sequencing and data analysis

Strand specific RNA sequencing libraries were constructed using the Illumina TruSeq RNA Access Library Prep Kit (Illumina, San Diego, CA) according to the manufacturer’s protocol. Briefly, the double stranded cDNA was hybridized to biotinylated, coding RNA capture probes. The resulting transcriptome-enriched library was sequenced by an Illumina HiSeq4000 using the 76 base pair paired end configuration. Raw sequencing data from the Illumina platform were converted to fastq files and aligned to the reference genome (hg19) using the STAR algorithm in single-pass mode with default parameters ^34^. HTSeq-count was then utilized to generate the raw counts for each gene^35^. Raw counts were then analyzed by DESeq2 for data processing, normalization and differential expression analysis according to standard procedures^36^.

### Single-cell targeted DNA sequencing and data analysis

We performed a single-cell targeted DNA sequencing using Tapestri^®^ platform (Mission Bio, South San Francisco, CA) as previously described ^37^. Briefly, frozen bone marrow cells were thawed and resuspended with lysis buffer. Each cell was encapsulated into the microfluidic droplet, then was barcoded to label each cell differently. Barcoded samples were amplified using 50 primer pairs specific to the 19 mutated AML genes covering known disease-related hotspot loci (Table S3). The pooled library was sequenced on an Illumina Miseq with 150-basepair paired end multiplexed runs. Fastq files generated from the MiSeq machine were processed using the Tapestri Analysis Pipeline (https://support.missionbio.com/hc/en-us/categories/360002512933-Tapestri-DNA-Pipeline) for adapter trimming, sequence alignment, barcode demultiplexing, and genotype and variant calling. Loom files generated by the pipeline were then analyzed by the in-house pipeline for variant annotation, filtering and results visualization.

### NetBID activity analysis

We performed NetBID2 analysis to identify hidden drivers of methylation-based Cluster 1 and Cluster 2 using RNA-seq data of baseline samples. We used normalized Log2 read count from RNA-seq of 27 Cluster 1 and 13 Custer 2 samples as input to generate networks using SJARACNe^38^. We used P value < 0.01 and log FC > 0.1 to select drivers.

### LSC17 score calculation

LSC17 score was calculated using RNA-seq data from baseline samples according to the equation published previously^23^: LSC17 score = (*DNMT3B* × 0.0874) + (*ZBTB46* × −0.0347) + (*NYNRIN* × 0.00865) + (*ARHGAP22* × −0.0138) + (*LAPTM4B* × 0.00582) + (*MMRN1* × 0.0258) + (*DPYSL3* × 0.0284) + (*KIAA0125* × 0.0196) + (*CDK6* × −0.0704) + (*CPXM1* × −0.0258) + (*SOCS2* × 0.0271) + (*SMIM24* × −0.0226) + (*EMP1* × 0.0146) + (*NGFRAP1* × 0.0465) + (*CD34* × 0.0338) + (*AKR1C3* × −0.0402) + (*GPR56* × 0.0501).

### Statistical analysis

The Chi-square or Fisher’s exact test was used to assess statistical differences in categorical variables and odds ratio to evaluate the strength of association. The Mann-Whitney U test or Student *t* test was used to analyze differences in continuous variables. Multiple hypothesis testing was corrected by Benjamini-Hochberg method. ROC curve as well as AUROC value were generated by pROC R package. A multivariate logistic regression model was performed to examine the relationship between CR variable and the predictors of LSC17 Score, RUNX1 mutation, RAS/RTK mutation and ELN high risk cytogenetics. All applicable tests were twosided and p value less than 0.05 was considered as statistical significance. Statistical analyses were performed within the analytic software described above or by R computing software (ver. 3.3.2).

## Data availability

Raw methylation and RNA-Seq data was submitted to GEO with the accession number GSE153349. The full list of detected baseline driver mutations was shown in Table S4.

## Code availability

The custom codes that support the findings of this study is available in GitHub (https://github.com/farmerkingwf/IDH_codes.git).

## Author contributions

Conception and design of the study was performed by K.T., C.D.D., and P.A.F.. K.T. and P.A.F. supervised the study and provided financial support. C.D.D. and H.K. led all the clinical trials relevant to this study. Sample collection and data assembly were performed by F.K., K.P., K.MB., B.W., G.L., M.F., J.M., L.L., C.G., T.K., G.GM., E.J., F.R., M.K., K.B., H.K., and K.T. F.W., K.M., S.X., J.Z., and K.T. performed data analysis and interpretation. F.W., and K.T. wrote the manuscript and revised based on the input from all other authors. All authors approved the final version of the manuscript.

## Acknowledgement

This study was supported in part by the Cancer Prevention Research Institute of Texas (grant R120501 to PAF), the Welch Foundation (grant G-0040 to PAF), the UT System STARS Award (grant PS100149 to PAF), Khalifa Scholar Award (to KT), Physician Scientist Program at MD Anderson (to KT), V Foundation Lloyd Family Clinical Scholar Award (to CDD), Charif Souki Cancer Research Fund (to HK), MD Anderson Cancer Center Leukemia SPORE P50 CA100632 (to HK), MD Anderson Cancer Center Support Grant (NIH P30 CA016672), Lynda Hill Foundation (PAF), JSPS Overseas Research Fellowship (TT), and by generous philanthropic contributions to MD Anderson’s Moon Shot Program for AML/MDS Platform (to HK, GGM and KT). We thank Sunita Patterson at Department Scientific Publications at MD Anderson for providing scientific editing of the manuscript.

## Reference

1 Papaemmanuil, E. et al. Genomic Classification and Prognosis in Acute Myeloid Leukemia. New England Journal of Medicine 374, 2209–2221, doi:10.1056/NEJMoa1516192 (2016).

2 Dang, L. et al. Cancer-associated IDH1 mutations produce 2-hydroxyglutarate. Nature 462, 739–744, doi: 10.1038/nature08617 (2009).

3 Ward, P. S. et al. The common feature of leukemia-associated IDH1 and IDH2 mutations is a neomorphic enzyme activity converting alpha-ketoglutarate to 2-hydroxyglutarate. Cancer cell 17, 225–234, doi:10.1016/j.ccr.2010.01.020 (2010).

4 Xu, W. et al. Oncometabolite 2-hydroxyglutarate is a competitive inhibitor of alphaketoglutarate-dependent dioxygenases. Cancer cell 19, 17–30, doi: 10.1016/j.ccr.2010.12.014 (2011).

5 Losman, J. A. et al. (R)-2-hydroxyglutarate is sufficient to promote leukemogenesis and its effects are reversible. Science 339, 1621–1625, doi:10.1126/science.1231677 (2013).

6 Zhao, S. et al. Glioma-derived mutations in IDH1 dominantly inhibit IDH1 catalytic activity and induce HIF-1alpha. Science 324, 261–265, doi:10.1126/science.1170944 (2009).

7 Lu, C. et al. IDH mutation impairs histone demethylation and results in a block to cell differentiation. Nature 483, 474–478, doi:10.1038/nature10860 (2012).

8 Figueroa, M. E. et al. Leukemic IDH1 and IDH2 mutations result in a hypermethylation phenotype, disrupt TET2 function, and impair hematopoietic differentiation. Cancer cell 18, 553–567, doi:10.1016/j.ccr.2010.11.015 (2010).

9 Amatangelo, M. D. et al. Enasidenib induces acute myeloid leukemia cell differentiation to promote clinical response. Blood, doi:10.1182/blood-2017-04-779447 (2017).

10 DiNardo, C. D. et al. Durable Remissions with Ivosidenib in IDH1-Mutated Relapsed or Refractory AML. N Engl J Med 378, 2386–2398, doi:10.1056/NEJMoa1716984 (2018).

11 Stein, E. M. et al. Enasidenib in mutant-<em>IDH2</em> relapsed or refractory acute myeloid leukemia. Blood, doi:10.1182/blood-2017-04-779405 (2017).

12 Wang, F. et al. Targeted inhibition of mutant IDH2 in leukemia cells induces cellular differentiation. Science 340, 622–626, doi:10.1126/science.1234769 (2013).

13 Kernytsky, A. et al. IDH2 mutation-induced histone and DNA hypermethylation is progressively reversed by small-molecule inhibition. Blood 125, 296–303, doi: 10.1182/blood-2013-10-533604 (2015).

14 Yen, K. et al. AG-221, a First-in-Class Therapy Targeting Acute Myeloid Leukemia Harboring Oncogenic IDH2 Mutations. Cancer Discov 7, 478–493, doi: 10.1158/2159-8290.CD-16-1034 (2017).

15 Quek, L. et al. Clonal heterogeneity of acute myeloid leukemia treated with the IDH2 inhibitor enasidenib. Nature Medicine, doi: 10.1038/s41591-018-0115-6 (2018).

16 Intlekofer, A. M. et al. Acquired resistance to IDH inhibition through trans or cis dimer-interface mutations. Nature 559, 125–129, doi:10.1038/s41586-018-0251-7 (2018).

17 Harding, J. J. et al. Isoform Switching as a Mechanism of Acquired Resistance to Mutant Isocitrate Dehydrogenase Inhibition. Cancer Discovery, doi:10.1158/2159-8290.Cd-18-0877 (2018).

18 Glass, J. L. et al. Epigenetic Identity in AML Depends on Disruption of Nonpromoter Regulatory Elements and Is Affected by Antagonistic Effects of Mutations in Epigenetic Modifiers. Cancer Discov 7, 868–883, doi:10.1158/2159-8290.CD-16-1032 (2017).

19 Du, X. et al. Hippo/Mst signalling couples metabolic state and immune function of CD8alpha(+) dendritic cells. Nature 558, 141–145, doi:10.1038/s41586-018-0177-0 (2018).

20 Adane, B. et al. The Hematopoietic Oxidase NOX2 Regulates Self-Renewal of Leukemic Stem Cells. Cell Reports 27, 238–254.e236, doi:https://doi.org/10.1016/j.celrep.2019.03.009 (2019).

21 Scheicher, R. et al. CDK6 as a key regulator of hematopoietic and leukemic stem cell activation. Blood 125, 90–101, doi:10.1182/blood-2014-06-584417 (2015).

22 Chung, S. S. et al. CD99 is a therapeutic target on disease stem cells in myeloid malignancies. Sci Transl Med 9, doi: 10.1126/scitranslmed.aaj2025 (2017).

23 Ng, S. W. K. et al. A 17-gene stemness score for rapid determination of risk in acute leukaemia. Nature 540, 433–437, doi:10.1038/nature20598 (2016).

24 Quek, L. et al. Clonal heterogeneity of acute myeloid leukemia treated with the IDH2 inhibitor enasidenib. Nat Med 24, 1167–1177, doi:10.1038/s41591-018-0115-6 (2018).

25 Choe, S. et al. Molecular mechanisms mediating relapse following ivosidenib monotherapy in IDH1-mutant relapsed or refractory AML. Blood Adv 4, 1894–1905, doi: 10.1182/bloodadvances.2020001503 (2020).

26 Grossmann, V. et al. Whole-exome sequencing identifies somatic mutations of BCOR in acute myeloid leukemia with normal karyotype. Blood 118, 6153–6163, doi: 10.1182/blood-2011-07-365320 (2011).

27 Cheson, B. D. et al. Revised recommendations of the International Working Group for Diagnosis, Standardization of Response Criteria, Treatment Outcomes, and Reporting Standards for Therapeutic Trials in Acute Myeloid Leukemia. J Clin Oncol 21, 4642–4649, doi:10.1200/JCO.2003.04.036 (2003).

28 Takahashi, K. et al. Preleukaemic clonal haemopoiesis and risk of therapy-related myeloid neoplasms: a case-control study. Lancet Oncol 18, 100–111, doi:10.1016/S1470-2045(16)30626-X (2017).

29 Takahashi, K. et al. Integrative genomic analysis of adult mixed phenotype acute leukemia delineates lineage associated molecular subtypes. Nat Commun 9, 2670, doi: 10.1038/s41467-018-04924-z (2018).

30 Morris, T. J. et al. ChAMP: 450k Chip Analysis Methylation Pipeline. Bioinformatics 30, 428–430, doi:10.1093/bioinformatics/btt684 (2014).

31 Zhou, W., Laird, P. W. & Shen, H. Comprehensive characterization, annotation and innovative use of Infinium DNA methylation BeadChip probes. Nucleic acids research 45, e22, doi:10.1093/nar/gkw967 (2017).

32 Teschendorff, A. E. et al. A beta-mixture quantile normalization method for correcting probe design bias in Illumina Infinium 450 k DNA methylation data. Bioinformatics 29, 189–196, doi:10.1093/bioinformatics/bts680 (2013).

33 Smyth, G. K. Linear models and empirical bayes methods for assessing differential expression in microarray experiments. Stat Appl Genet Mol Biol 3, Article3, doi: 10.2202/1544-6115.1027 (2004).

34 Dobin, A. et al. STAR: ultrafast universal RNA-seq aligner. Bioinformatics 29, 15–21, doi: 10.1093/bioinformatics/bts635 (2013).

35 Anders, S., Pyl, P. T. & Huber, W. HTSeq—a Python framework to work with high-throughput sequencing data. Bioinformatics 31, 166–169, doi: 10.1093/bioinformatics/btu638 (2015).

36 Love, M. I., Huber, W. & Anders, S. Moderated estimation of fold change and dispersion for RNA-seq data with DESeq2. Genome Biology 15, 550, doi: 10.1186/s13059-014-0550-8 (2014).

37 Pellegrino, M. et al. High-throughput single-cell DNA sequencing of acute myeloid leukemia tumors with droplet microfluidics. Genome Res 28, 1345–1352, doi: 10.1101/gr.232272.117 (2018).

38 Khatamian, A., Paull, E. O., Califano, A. & Yu, J. SJARACNe: a scalable software tool for gene network reverse engineering from big data. Bioinformatics 35, 2165–2166, doi: 10.1093/bioinformatics/bty907 (2019).

