## Supplementary Materials for "Leukemia stemness and co-occurring mutations drive resistance to IDH inhibitors in acute myeloid leukemia"

#### Contents

**Figure S1.**
**Figure S2.**
**Figure S3.**
**Figure S4.**
**Figure S5.**
**Figure S6.**
**Figure S7.**
**Figure S8.**
**Figure S9.**
**Figure S10.**
**Figure S11.**
**Figure S12.**
**Figure S13.**
**Figure S14.**
**Table S1.**
**Table S2.**
**Table S3.**
**Table S4.**
**Supplemental Methods.**

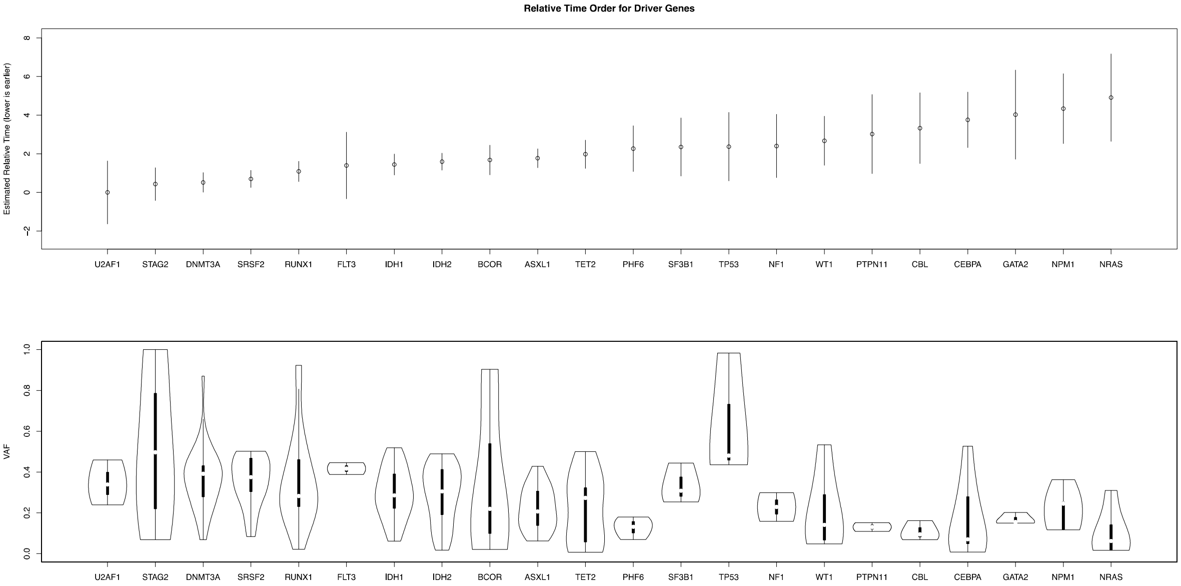


**Figure S1. Relative timing of the mutation accrual in the IDHi-treated AML patients.** Forrest plots (top) showing the relative timing of mutation accrual for each driver genes detected in the baseline samples of the IDHi-treated AML patients. The dots represent the ability estimates based on an unstructured Bradley-Terry model. The lower the ability estimates, the earlier the relative timing would be. The error bars represent the standard errors. Violin plots (bottom) showing the driver mutation VAF distributions for all driver genes.

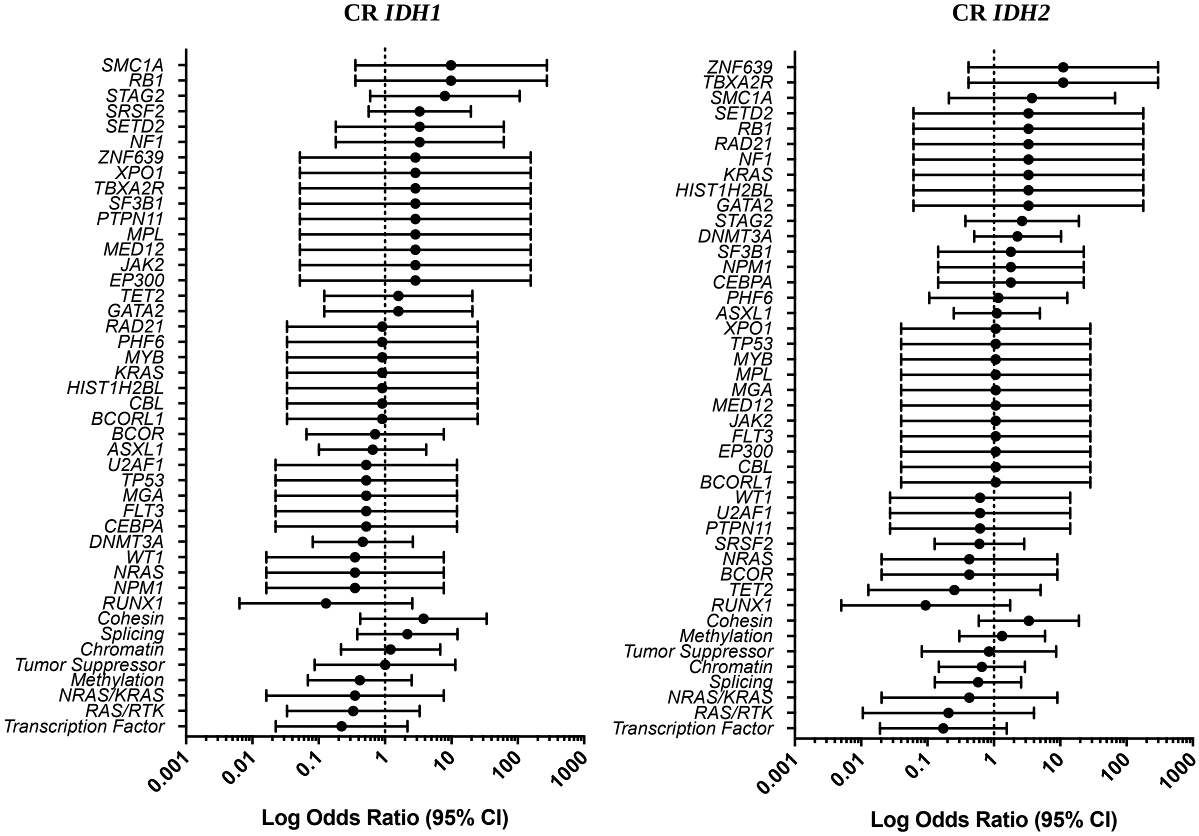


**Figure S2. Forrest plots showing enrichment of the mutations at baseline against Complete Remission (CR) by logarithmic odds ratio for *IDH1*- (left) and *IDH2*- (right) mutated patients.** *$P < 0.05$. Circles represent odds ratios. The error bars represent 95% confidence interval of odds ratio.

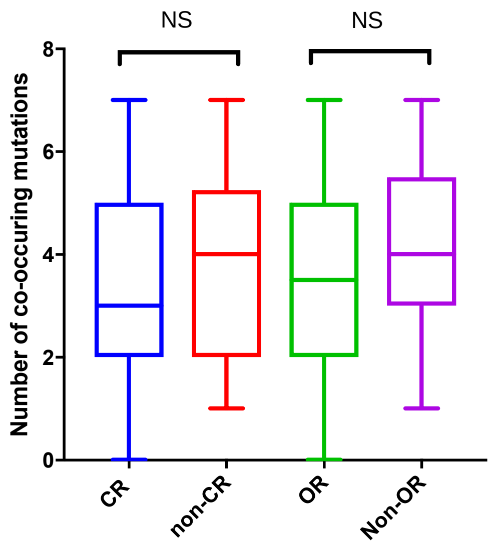


**Figure S3. The number of co-occurring mutations at baseline did not have a significant impact with respect to the clinical response in our cohort.** Box plots comparing the number of co-occurring mutations in baseline samples from patients who achieved CR, non-CR, OR or non-OR. Abbreviations: CR, complete remission; OR, overall response; NS, non-significance.

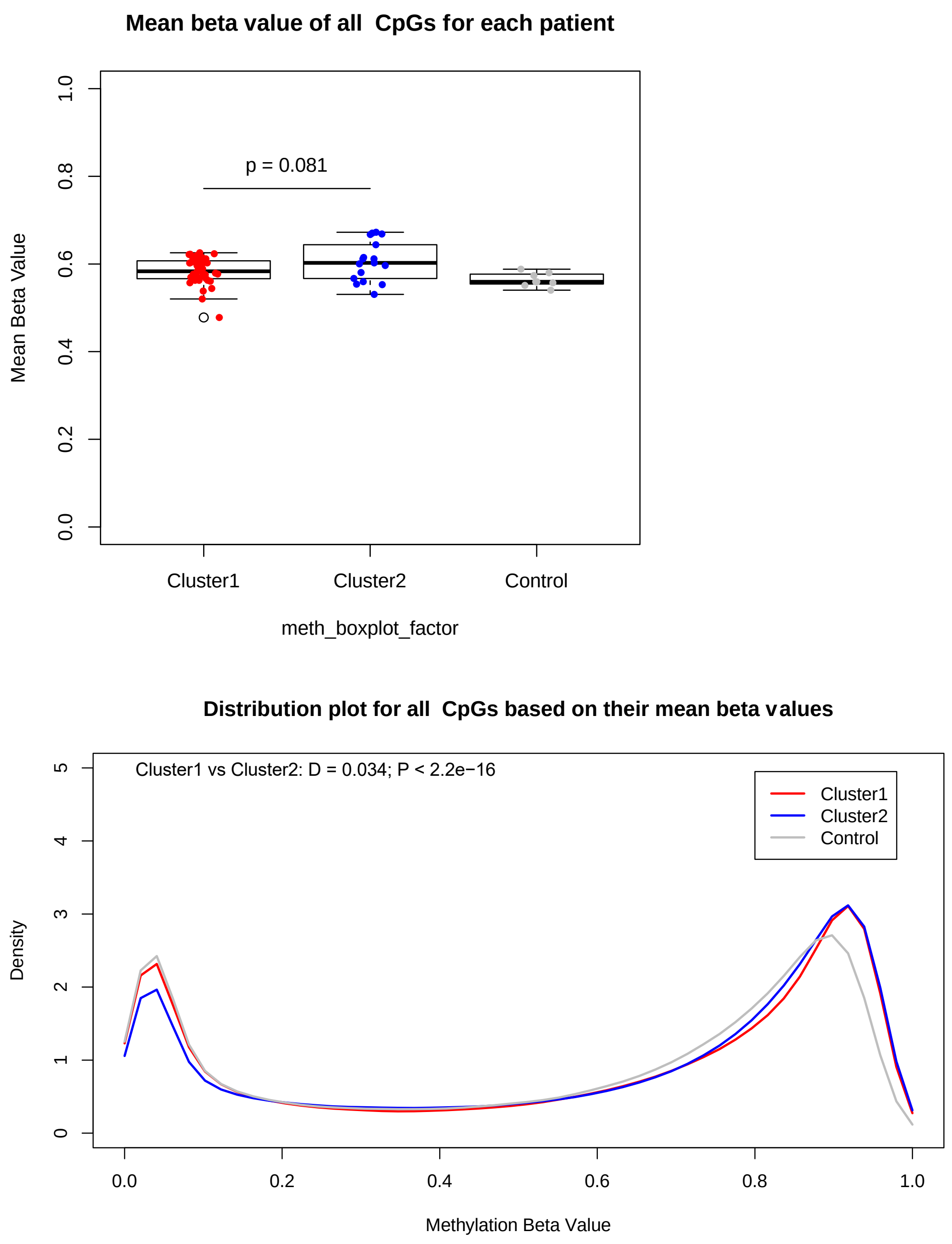


**Figure S4. Cluster 2 patients show hypermethylation at baseline time point when compared to Cluster 1 patients.** (Top) Box plot comparing mean methylation beta value of all CpGs among Cluster 1 baseline (N=40) and Cluster 2 baseline (N=17) samples. IDH1/2 wild type AML samples (N=8) are used as control. (Bottom) Density distribution of all CpG probes with methylation beta values comparing Cluster 1 baseline and Cluster 2 samples. Kolmogorov–Smirnov test D and P values are shown. IDH1/2 wild type AML samples (N=8) are used as control.

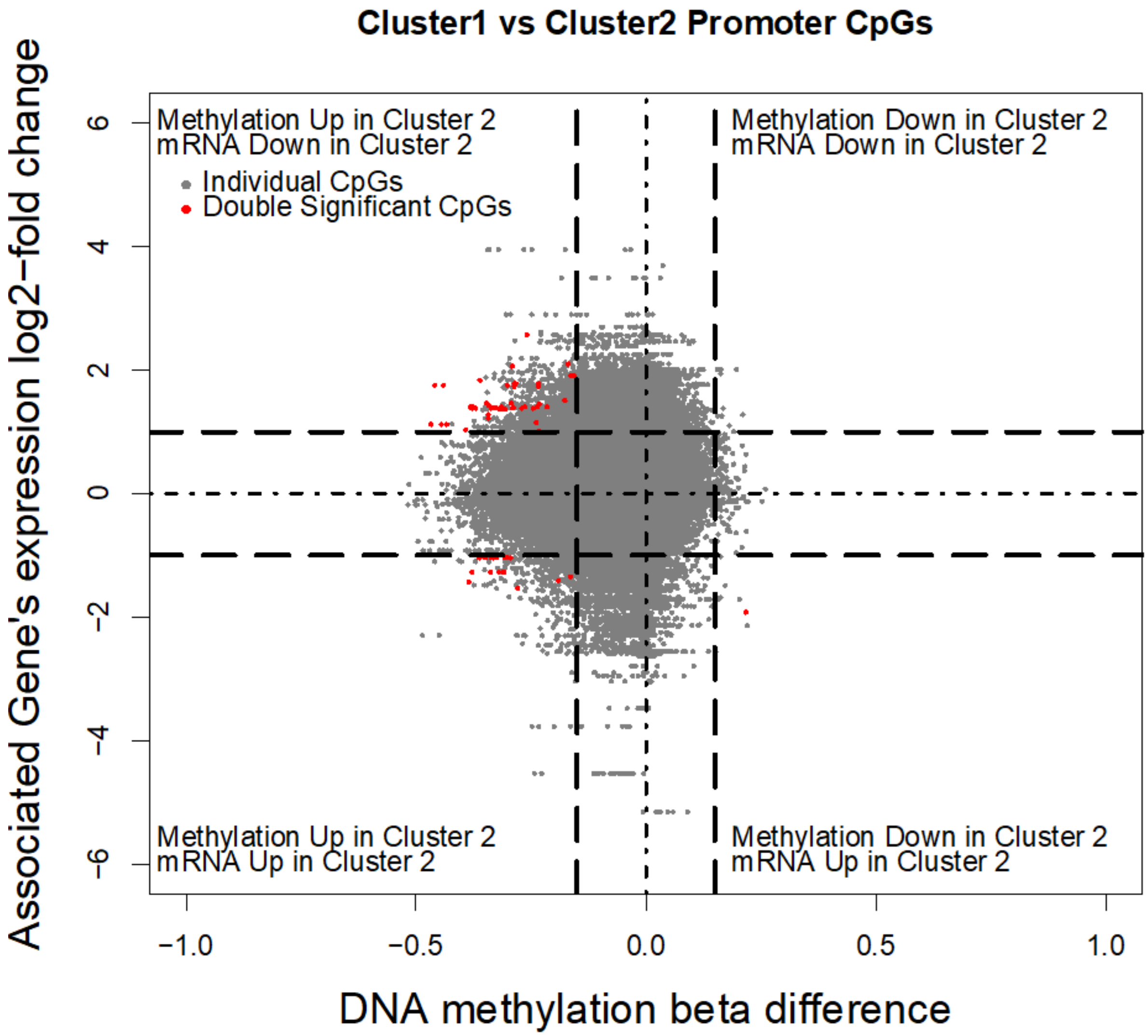


**Figure S5. Starburst plot showing integrated analysis of gene expression and promoter methylation changes between Cluster 1 and Cluster 2.** Of 215521 promoter CpGs, 67 showed statistically significant differential methylation and differential expression between Cluster 1 and Cluster 2. More than half of them showed promoter hypermethylation and downregulation of the gene expression.

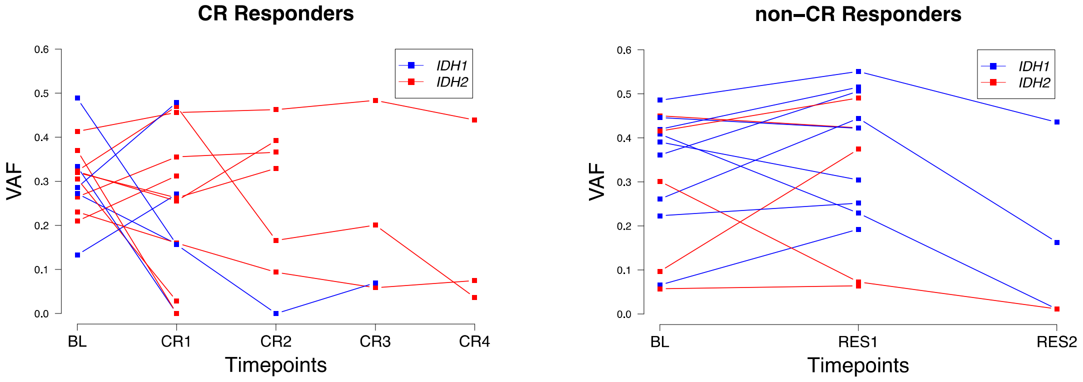


**Figure S6. Longitudinal trend of VAF for *IDH1/2* mutations for responders achieving CR (left) and all other non-CR (right) responders.** Abbreviations: CR, complete remission; VAF, variant allele frequency; BL, baseline; RES, response.

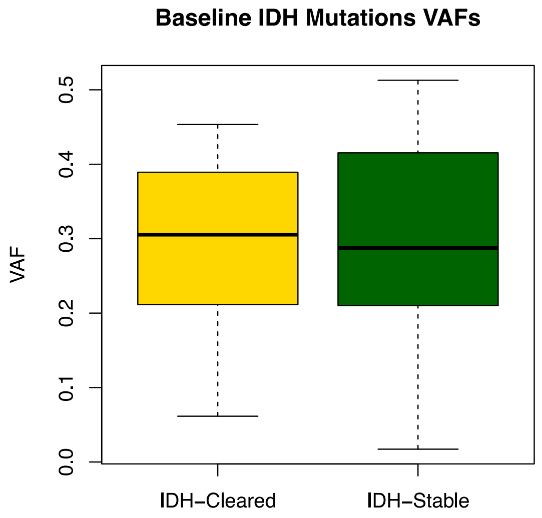


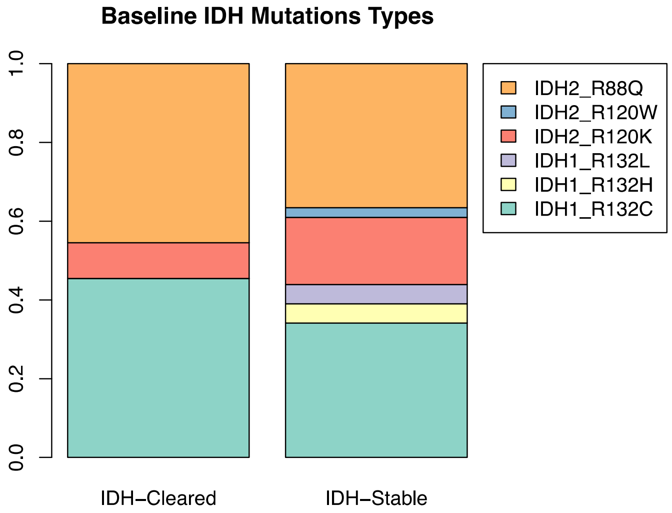


**Figure S7. The baseline VAF or the types of *IDH* mutations did not predict the clearance of the mutations.** Box plots (top) showing the VAF distribution of *IDH* mutations in *IDH*-Cleared and *IDH*-Stable baseline samples. Bar plots (bottom) showing the mutation-type distribution in *IDH*-Cleared and *IDH*-Stable baseline samples.

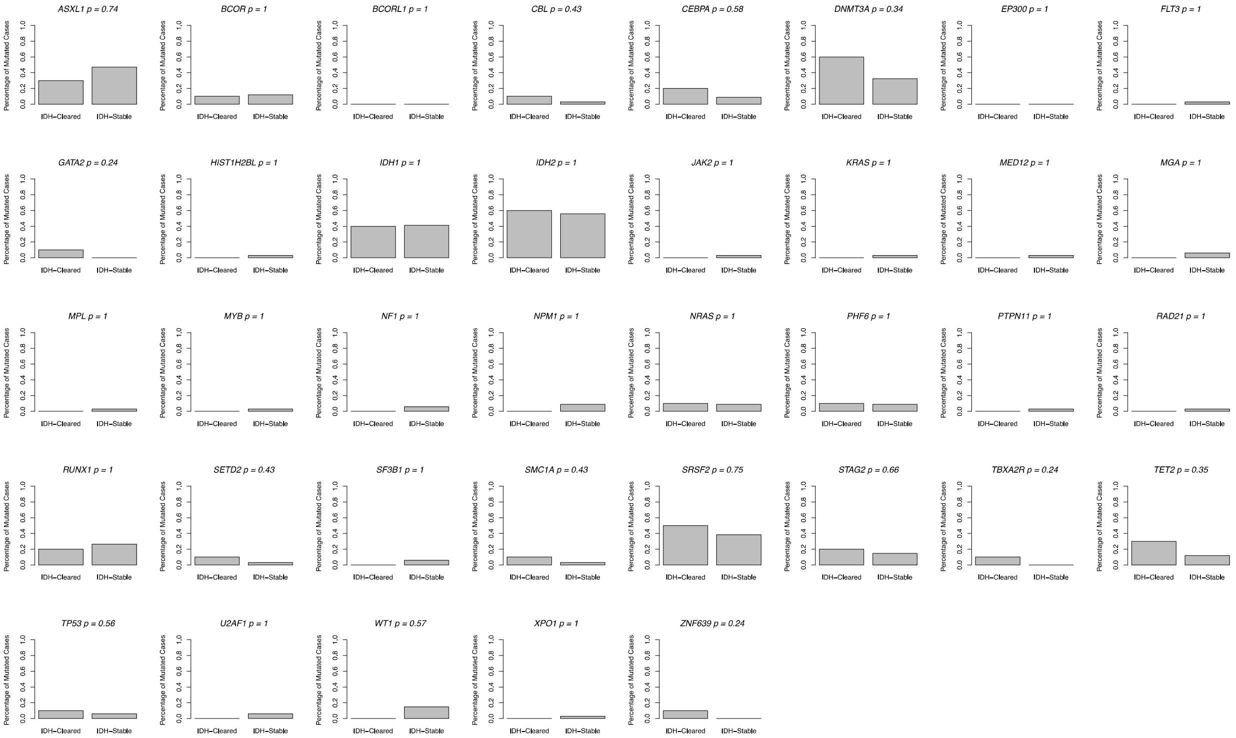


**Figure S8. No correlation between *IDH* mutation clearance and the co-occurring mutations.** Bar plots showing the mutated case percentage in *IDH*-Cleared and *IDH*-Stable baseline samples for each co-mutated gene.

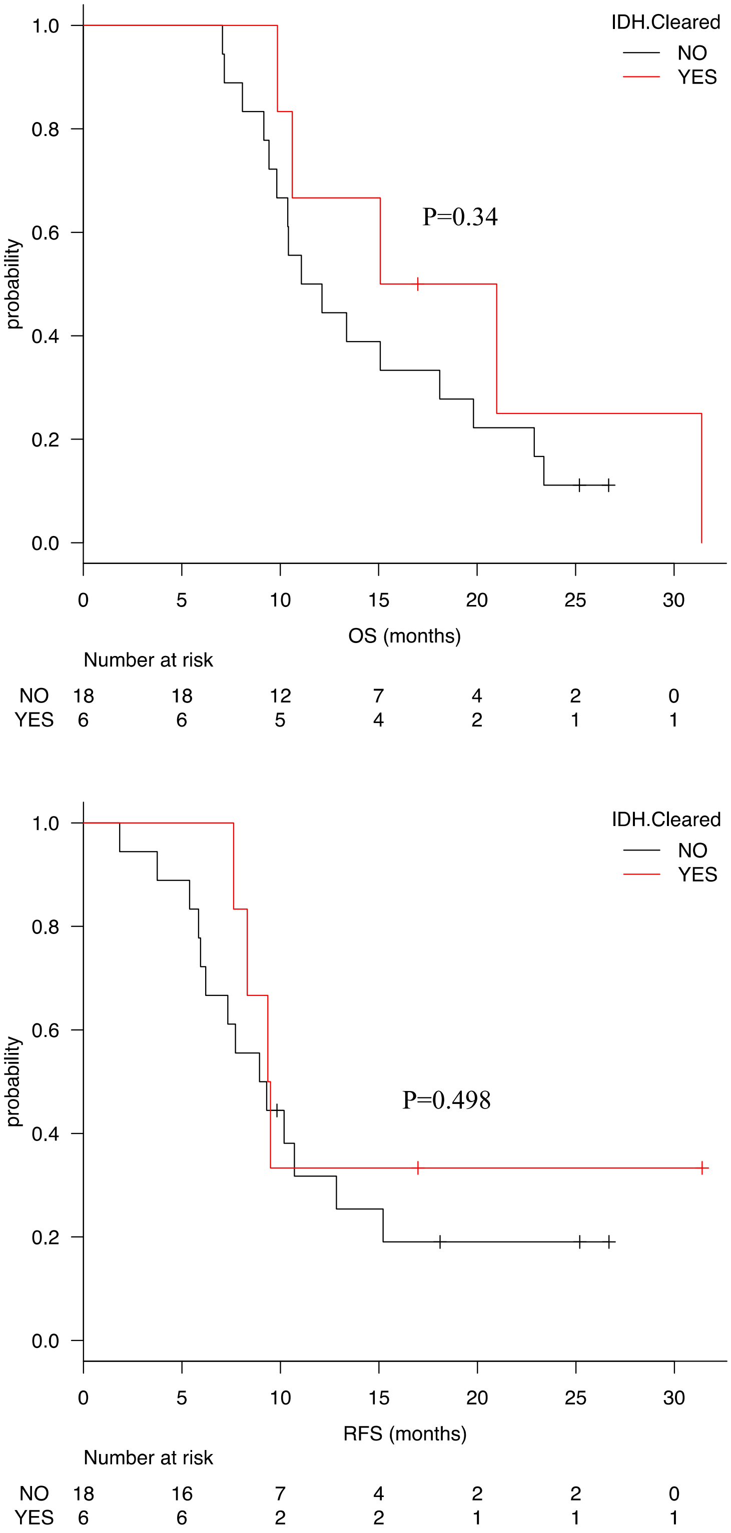


**Figure S9. Overall survival (OS, top) and Relapse-free survival (RFS, bottom) comparing *IDH*-Cleared and *IDH*-Stable patients.**

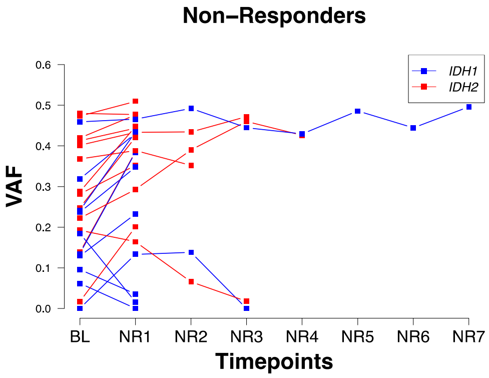


**Figure S10. Longitudinal trend of VAF for *IDH1/2* mutations for non-responders.**
Abbreviations: VAF, variant allele frequency; BL, baseline; NR, non-response.

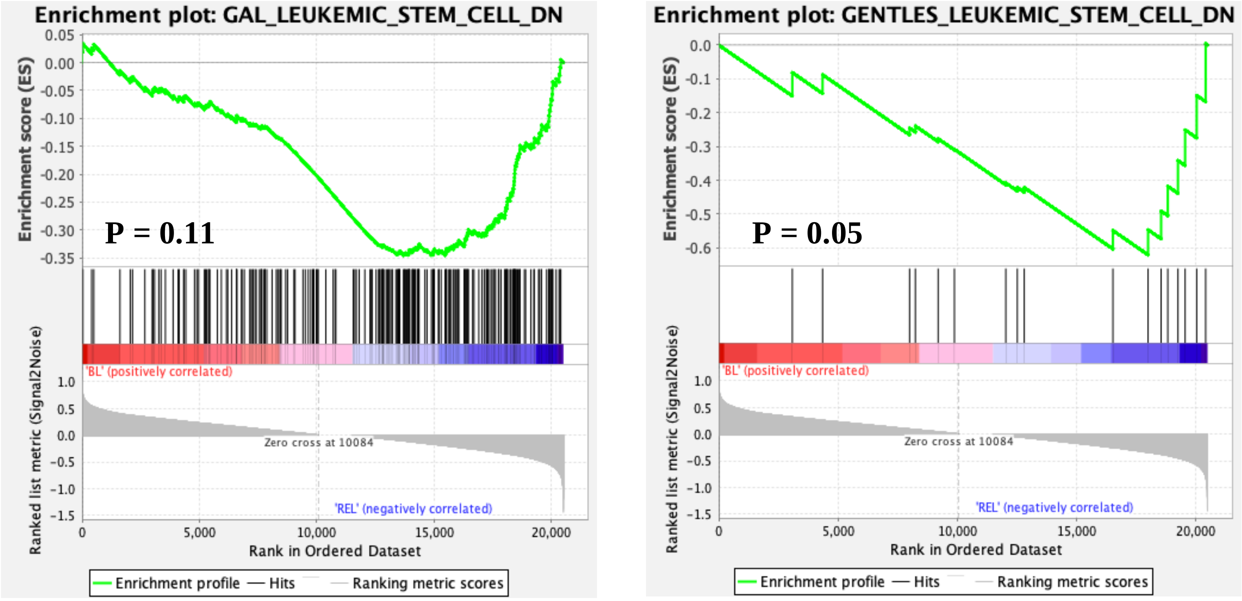


**Figure S11. Gene Set Enrichment Analysis comparing gene expression profiles between baseline and relapse samples.** Genes downregulated in leukemia stem cells (LSC) are enriched in relapse samples.

UPI2370759

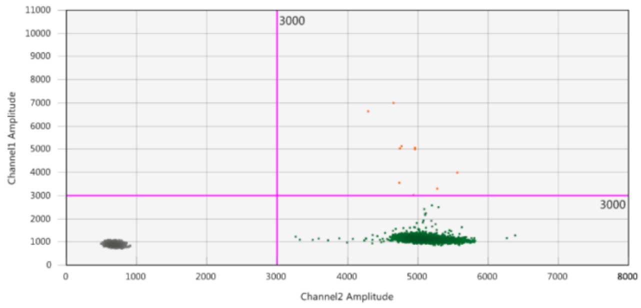


Positive control

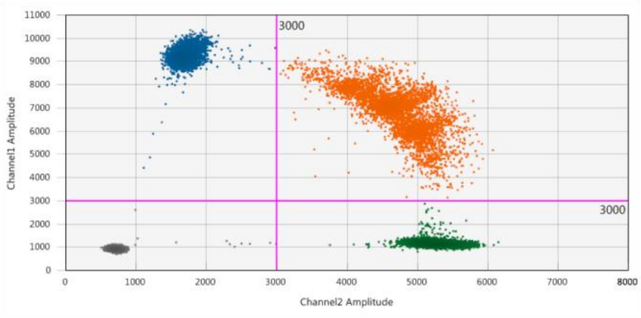


Negative control

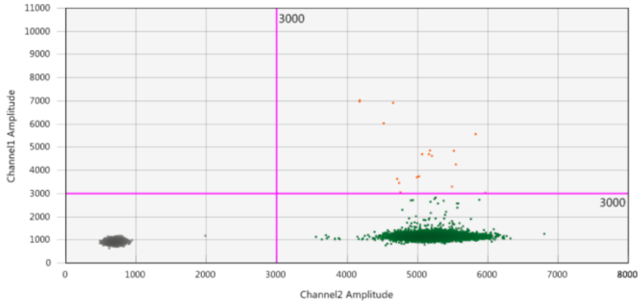


No template control

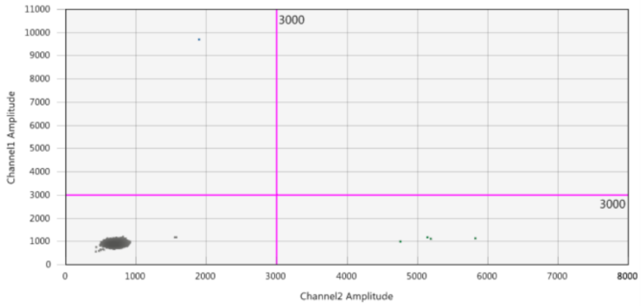


**Figure S12. ddPCR did not detect *IDH1* R132C in UPI2370759.** 2-D plots showing amplitude in two channels. Blue cluster shown in upper-left quadrant represents droplets with mutant DNA only. Orange cluster shown in upper-right quadrant represents droplets with both mutant and wildtype DNA. Green cluster shown in lower-right quadrant represents droplets with wildtype DNA only. Grey cluster shown in lower-left quadrant represents droplets without DNA from targeted locus. Fractional abundance was 0.0714% in UPI2370759.

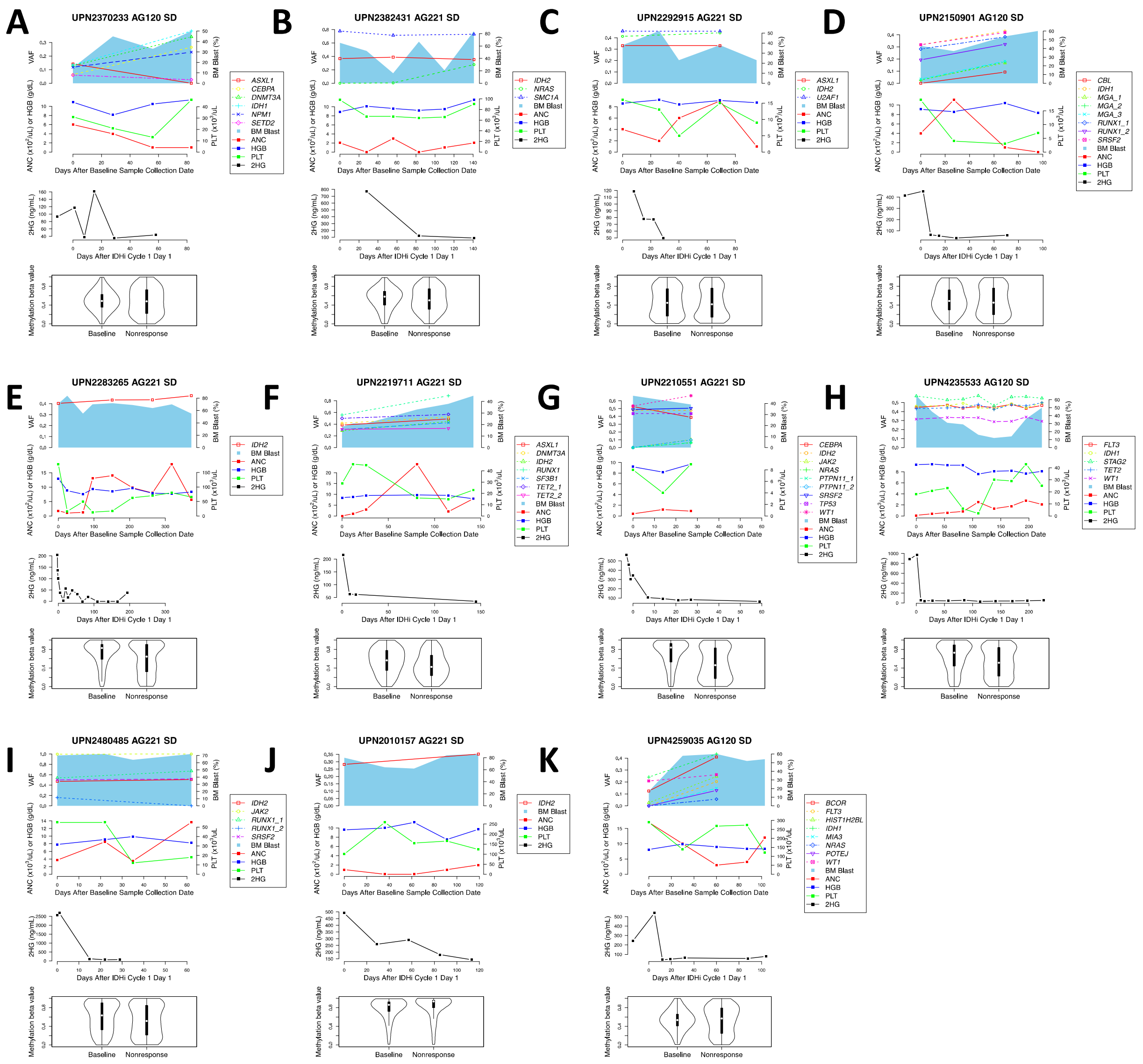

**Figure S13. Genetic and epigenetic co-evolution in 11 non-responders.** (**A-I**) Cases that show co-suppression of 2HG and DNA methylation after IDH inhibitor. (**J-K**) Cases that show 2HG suppression but without demethylation.

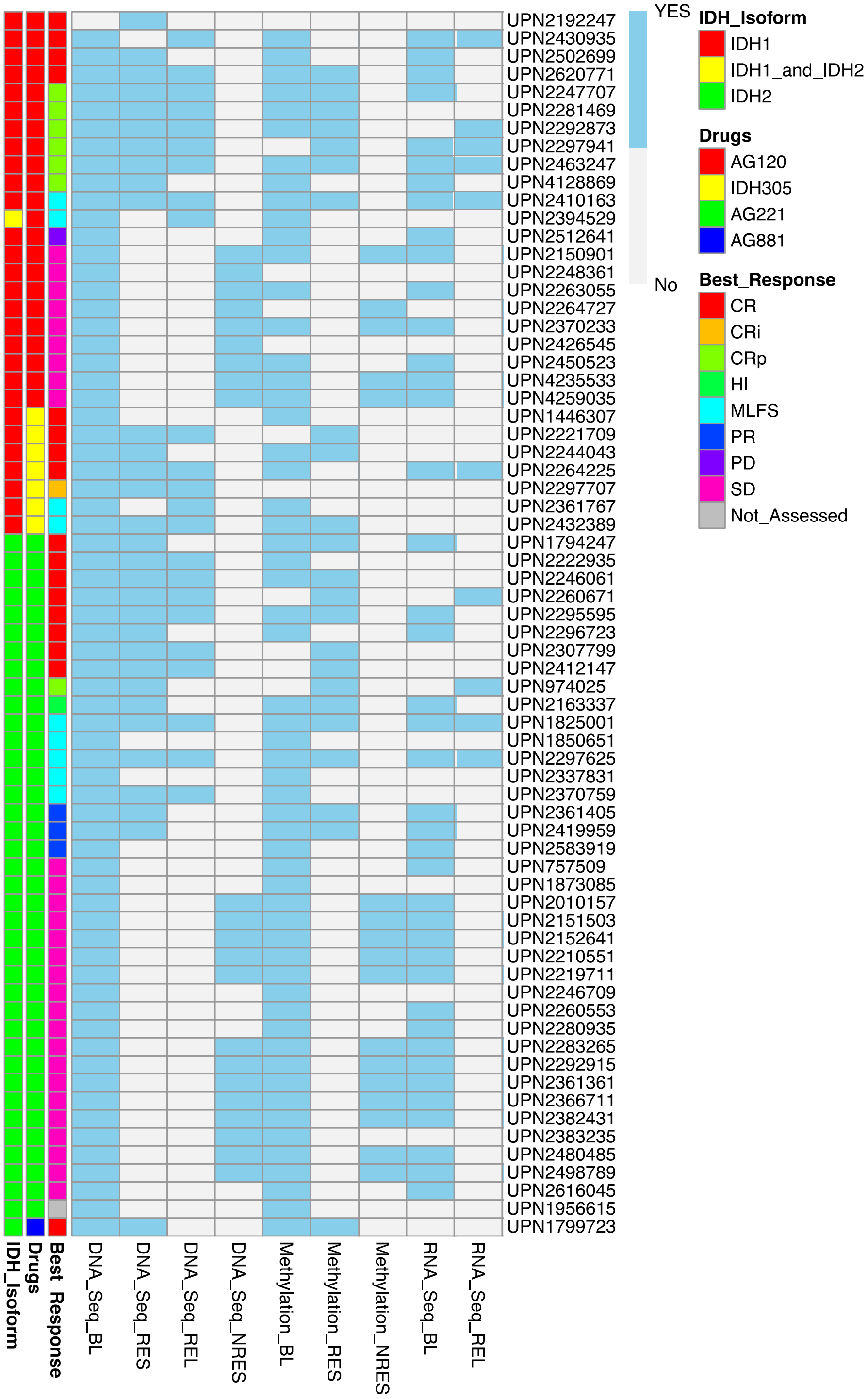


**Figure S14. Summary heatmap for sample availability across all analytic platforms for all 68 patients.** Abbreviations: BL, baseline; RES, response; REL, relapse; NRES, non-response.

**Table S1.** Comparison of clinical features between included and excluded patients.

| | included (N=68) | | excluded (N=80) | | |
|---|---|---|---|---|---|
| | **median** | **IQR** | **median** | **IQR** | p |
| **WBC** | 1.5 | 0.9-2.3 | 1.8 | 1.0-8.0 | 0.084 |
| **ANC** | 0.2 | 0.1-0.6 | 0.5 | 0.2-1.7 | 0.006 |
| **HGB** | 9.4 | 9.8-10.1 | 9.4 | 8.8-10.2 | 0.965 |
| **PLT** | 33 | 20-64 | 35 | 19-71 | 0.913 |
| **BM blast, %** | 39 | 18-63 | 27 | 9-56 | 0.045 |
| **Age** | 72 | 65-77 | 67 | 58-73 | 0.01 |
| | **No.** | **%** | **No.** | **%** | |
| **Karyotype** | | | | | 1 |
| intermediate | 46 | 70 | 55 | 70 | |
| poor | 20 | 30 | 24 | 30 | |
| **Sex** | | | | | 0.498 |
| female | 28 | 41 | 28 | 35 | |
| male | 40 | 59 | 52 | 65 | |
| **Drug** | | | | | <0.001 |
| AG120 | 22 | 32 | 33 | 41 | |
| AG221 | 38 | 56 | 18 | 23 | |
| AG881 | 1 | 2 | 17 | 21 | |
| IDH305 | 7 | 10 | 12 | 15 | |
| **Best Response** | | | | | |
| CR | 17 | 25 | 5 | 6 | |
| CRi | 1 | 2 | 1 | 1 | |
| CRp | 7 | 10 | 4 | 5 | |
| MLFS | 9 | 13 | 5 | 6 | |
| PR | 3 | 4 | 2 | 3 | |
| HI | 1 | 2 | 2 | 3 | |
| SD | 28 | 41 | 29 | 36 | |
| PD | 1 | 2 | 18 | 23 | |
| Not done | 1 | 2 | 14 | 18 | |
| Responder | 38 | 57 | 19 | 29 | 0.002 |
| Non-Responder | 29 | 43 | 47 | 71 | |

ND – Student's t-test; non-ND – Mann–Whitney U test

WBC – non-ND; ANC– non-ND; HGB – ND; PLT– non-ND; BM blast, %– non-ND; Age– non-ND

Abbreviations: ND, normal distribution

#### Table S2. List of 295 genes targeted by next generation sequencing.

| | | | | | | | | | |
|---|---|---|---|---|---|---|---|---|---|
| *ABCC9* | *CALR* | *CUL5* | *FANCD2* | *HIST1H2BF* | *LEF1* | *NBN* | *PLA2G2D* | *SF3B1* | *TINF2 (TIN2)* |
| *ABL1* | *CARD11* | *CUX1* | *FANCE* | *HIST1H3D* | *LRP1B* | *NCOR1* | *PLCG2* | *SFRS1* | *TLR2* |
| *ACTG1* | *CBL* | *CYLD* | *FANCG* | *HIST1H4D* | *LTB* | *NCOR2* | *POT1* | *SFRS7* | *TLR9* |
| *AKT1* | *CBLB* | *DAXX* | *FANCI* | *HNRNPK* | *LUC7L2* | *NF1* | *POU2AF1* | *SGK1* | *TNFAIP3* |
| *ANKRD11* | *CCND1* | *DCLRE1C* | *FANCL* | *HRAS* | *LYN* | *NFE2* | *PRDM1* | *SH2B3* | *TNFRSF14* |
| *ARID1A* | *CCND3* | *DDX3X* | *FAS* | *ICOS* | *MALT1* | *NFKB1* | *PRKCB* | *SHH* | *TNKS* |
| *ARID1B* | *CD200* | *DIS3* | *FAT1* | *ID3* | *MAP2K1* | *NFKB2* | *PTEN* | *SMAD2* | *TOX* |
| *ARID2* | *CD274* | *DKC1* | *FAT3* | *IDH1* | *MAPK1* | *NFKBIA* | *PTPN1* | *SMC1A* | *TP53* |
| *ARID5B* | *CD58* | *DLC1* | *FBXW7* | *IDH2* | *MAX* | *NFKBIE* | *PTPN11* | *SMC3* | *TRAF3* |
| *ARPP21* | *CD79A* | *DNM2* | *FGFR3* | *IKBKA* | *MDM2* | *NOTCH1* | *RAD21* | *SMC5* | *TRAF6* |
| *ASXL1* | *CD79B* | *DNMT1* | *FLI1* | *IKZF1* | *MED12* | *NOTCH2* | *RAD51C* | *SNX7* | *TYK2* |
| *ATF7IP* | *CDK4* | *DNMT3A* | *FLT3* | *IKZF2* | *MEF2B* | *NPM1* | *RAG1* | *SOCS1* | *TYK3* |
| *ATM* | *CDKN2A* | *DNMT3B* | *FNDC3A* | *IKZF3* | *MEF2C* | *NR3C2* | *RAG2* | *SOX5* | *U2AF1* |
| *ATRX* | *CDKN2B* | *EBF1* | *FOXP1* | *IL7R* | *MGA* | *NRAS* | *RASA2* | *SP140* | *U2AF2* |
| *B2M* | *CDKN2C* | *ECT2L* | *FYN* | *IRAK1* | *miR125a* | *NSD2* | *RB1* | *SPEN* | *UBR5* |
| *BCL10* | *CEBPA* | *EED* | *G6PC3* | *IRAK4* | *miR-142* | *NT5C2* | *REL* | *SPIB* | *USP29* |
| *BCL2* | *CEBPE* | *EGR1* | *GAB2* | *IRF1* | *mIR155* | *PAG1* | *RELA* | *SRSF2* | *VPREB1* |
| *BCL6* | *CHD2* | *EGR2* | *GATA1* | *IRF4* | *mIR15a* | *PALB2* | *RELB* | *STAG1* | *WHSC1* |
| *BCL7A* | *CHK2* | *ELANE* | *GATA2* | *IRF7* | *mIR16-1* | *PAX5* | *RELN* | *STAG2* | *WHSC1L1* |
| *BCOR* | *CIITA* | *EP300* | *GATA3* | *ITPKB* | *MIR17HG* | *PDCD1* | *RHOA* | *STAT1* | *WT1* |
| *BCR* | *CNOT3* | *EPHA7* | *GCET2* | *JAK1* | *mIR21* | *PDCD1LG2* | *RIPK1* | *STAT3* | *XPO1* |
| *BIRC3* | *CREBBP* | *EPOR* | *GFI1B* | *JAK2* | *mir34b* | *PDGFRB* | *ROBO1* | *SUZ12* | *ZAP70* |
| *BLK* | *CRLF2* | *ERG* | *GNA13* | *JAK3* | *mir34c* | *PEG3* | *ROR1* | *SYK* | *ZMYM2* |
| *BMI1* | *CSF2RA* | *ETV6* | *GNAS* | *JARID2* | *MLL* | *PHF6* | *RPL10* | *TBL1XR1* | *ZMYM3* |
| *BRAF* | *CSF3R* | *EZH2* | *GNB1* | *KDM4C* | *MLL2* | *PHIP* | *RPL5* | *TCF3* | *ZRSR2* |
| *BRIP1* | *CTBP1* | *FAM46C* | *GPRC5A* | *KDM6A* | *MLL3* | *PIGA* | *RUNX1* | *TERC* | |
| *BTG1* | *CTBP2* | *FAM5C* | *HAX1* | *KIT* | *MPL* | *PIK3CA* | *RUNX2* | *TERT* | |
| *BTK* | *CTCF* | *FANCA* | *HIST1H1E* | *KLHL6* | *MS4A1* | *PIK3CB* | *SAMHD1* | *TET1* | |
| *BTLA* | *CTLA4* | *FANCB* | *HIST1H2AD* | *KRAS* | *MYB* | *PIK3CG* | *SETBP1* | *TET2* | |
| *C22orf194* | *CTNNA1* | *FANCC* | *HIST1H2BE* | *LAMB4* | *MYD88* | *PIK3R1* | *SETD2* | *TGDS* | |

**Table S3. List of 50 amplicons covered by the single cell DNA sequencing.**

| Amplicon | Chr | Primer Start | Insert Start | Insert End | Primer Start | Amplicon Length (without primer) [bp] | Amplicon Length (with primer) [bp] | Primer length [bp] |
|---|---|---|---|---|---|---|---|---|
| *ASXL1_1* | 20 | 31022348 | 31022369 | 31022586 | 31022608 | 217 | 260 | 43 |
| *ASXL1_2_a1* | 20 | 31022880 | 31022899 | 31023110 | 31023130 | 211 | 250 | 39 |
| *DNMT3A_10* | 2 | 25457114 | 25457135 | 25457351 | 25457372 | 216 | 258 | 42 |
| *EZH2_1* | 7 | 148504627 | 148504654 | 148504874 | 148504901 | 220 | 274 | 54 |
| *EZH2_2* | 7 | 148506303 | 148506332 | 148506547 | 148506577 | 215 | 274 | 59 |
| *FLT3_1* | 13 | 28592473 | 28592494 | 28592723 | 28592747 | 229 | 274 | 45 |
| *FLT3_2_a3* | 13 | 28608168 | 28608191 | 28608368 | 28608392 | 177 | 224 | 47 |
| *FLT3_3* | 13 | 28602155 | 28602179 | 28602404 | 28602429 | 225 | 274 | 49 |
| *FLT3_4* | 13 | 28609521 | 28609547 | 28609769 | 28609795 | 222 | 274 | 52 |
| *FLT3_5_4* | 13 | 28607997 | 28608018 | 28608153 | 28608176 | 135 | 179 | 44 |
| *GATA2_1* | 3 | 128202704 | 128202723 | 128202891 | 128202911 | 168 | 207 | 39 |
| *IDH1_1* | 2 | 209112875 | 209112898 | 209113123 | 209113149 | 225 | 274 | 49 |
| *IDH2_1_4* | 15 | 90631738 | 90631759 | 90631985 | 90632009 | 226 | 271 | 45 |
| *JAK2_1* | 9 | 5073541 | 5073563 | 5073785 | 5073815 | 222 | 274 | 52 |
| *KIT_1* | 4 | 55599204 | 55599231 | 55599448 | 55599478 | 217 | 274 | 57 |
| *KIT_2* | 4 | 55589585 | 55589607 | 55589829 | 55589859 | 222 | 274 | 52 |
| *KRAS_1* | 12 | 25398161 | 25398183 | 25398405 | 25398435 | 222 | 274 | 52 |
| *KRAS_2* | 12 | 25380238 | 25380259 | 25380466 | 25380490 | 207 | 252 | 45 |
| *NPM1_1_2* | 5 | 170837385 | 170837412 | 170837636 | 170837659 | 224 | 274 | 50 |
| *NRAS_1* | 1 | 115256296 | 115256324 | 115256546 | 115256570 | 222 | 274 | 52 |
| *NRAS_2* | 1 | 115258525 | 115258553 | 115258776 | 115258799 | 223 | 274 | 51 |
| *PTPN11_1_1* | 12 | 112926827 | 112926848 | 112927043 | 112927063 | 195 | 236 | 41 |
| *PTPN11_2* | 12 | 112888095 | 112888116 | 112888327 | 112888351 | 211 | 256 | 45 |
| *RUNX1_2* | 21 | 36171458 | 36171482 | 36171710 | 36171732 | 228 | 274 | 46 |
| *RUNX1_3* | 21 | 36206684 | 36206705 | 36206884 | 36206907 | 179 | 223 | 44 |
| *RUNX1_4* | 21 | 36231583 | 36231604 | 36231834 | 36231857 | 230 | 274 | 44 |
| *RUNX1_5* | 21 | 36164768 | 36164785 | 36164997 | 36165018 | 212 | 250 | 38 |
| *RUNX1_7* | 21 | 36252789 | 36252811 | 36253007 | 36253030 | 196 | 241 | 45 |
| *SF3B1_1* | 2 | 198266733 | 198266761 | 198266977 | 198267007 | 216 | 274 | 58 |
| *SF3B1_2* | 2 | 198267134 | 198267156 | 198267384 | 198267406 | 228 | 272 | 44 |
| *SRSF2_2_2* | 17 | 74732865 | 74732882 | 74733050 | 74733069 | 168 | 204 | 36 |
| *TP53_1* | 17 | 7578062 | 7578086 | 7578296 | 7578319 | 210 | 257 | 47 |
| *TP53_2* | 17 | 7577376 | 7577397 | 7577615 | 7577637 | 218 | 261 | 43 |
| *TP53_3* | 17 | 7578363 | 7578384 | 7578605 | 7578627 | 221 | 264 | 43 |
| *TP53_4* | 17 | 7576930 | 7576953 | 7577180 | 7577204 | 227 | 274 | 47 |

| | | | | | | | | |
|---|---|---|---|---|---|---|---|---|
| *U2AF1_1* | 21 | 44514679 | 44514700 | 44514922 | 44514947 | 222 | 268 | 46 |
| *U2AF1_2* | 21 | 44524258 | 44524279 | 44524505 | 44524532 | 226 | 274 | 48 |
| *WT1_1_a2* | 11 | 32414174 | 32414193 | 32414411 | 32414432 | 218 | 258 | 40 |
| *WT1_2* | 11 | 32413389 | 32413415 | 32413641 | 32413663 | 226 | 274 | 48 |
| *WT1_3* | 11 | 32417744 | 32417765 | 32417992 | 32418018 | 227 | 274 | 47 |
| chr10_106721610 | 10 | 106721487 | 106721508 | 106721711 | 106721736 | 203 | 249 | 46 |
| chr10_5554293 | 10 | 5554171 | 5554192 | 5554401 | 5554419 | 209 | 248 | 39 |
| chr10_77210191 | 10 | 77210064 | 77210083 | 77210294 | 77210313 | 211 | 249 | 38 |
| chr14_56969005 | 14 | 56968884 | 56968905 | 56969106 | 56969129 | 201 | 245 | 44 |
| chr16_55770629 | 16 | 55770512 | 55770529 | 55770735 | 55770757 | 206 | 245 | 39 |
| chr16_8569820 | 16 | 8569695 | 8569720 | 8569926 | 8569944 | 206 | 249 | 43 |
| chr18_9750662 | 18 | 9750543 | 9750561 | 9750767 | 9750791 | 206 | 248 | 42 |
| chr6_17076840 | 6 | 17076720 | 17076739 | 17076941 | 17076969 | 202 | 249 | 47 |
| chr6_40116264 | 6 | 40116143 | 40116164 | 40116366 | 40116388 | 202 | 245 | 43 |
| chr6_62094287 | 6 | 62094166 | 62094187 | 62094388 | 62094411 | 201 | 245 | 44 |

### Table S4. Driver mutation list of 67 baseline samples.

| UPN | GENE | TYPE | AA_CHANGE | VAF |
|---|---|---|---|---|
| **757509** | *DNMT3A* | nonsynonymous | DNMT3A:uc002rgd.4:exon17:c.C2012T:p.T671M | 0.508671 |
| **757509** | *EP300* | frameshift insertion | EP300:uc003azl.4:exon9:c.1771dupA:p.A590fs | 0.15254237 |
| **757509** | *IDH2* | nonsynonymous | IDH2:uc002box.3:exon4:c.C418T:p.R140W | 0.191589 |
| **757509** | *SRSF2* | nonframeshift deletion | SRSF2:uc010wtg.2:exon1:c.281_292del:p.94_98del | 0.25 |
| **757509** | *TET2* | frameshift deletion | TET2:uc021xqk.1:exon3:c.1363delC:p.P455fs | 0.05945946 |
| **974025** | *ASXL1* | frameshift deletion | ASXL1:uc021wbw.1:exon13:c.1926delA:p.G642fs | 0.162 |
| **974025** | *IDH2* | nonsynonymous | IDH2:uc002box.3:exon4:c.G515A:p.R172K | 0.45 |
| **974025** | *NRAS* | nonsynonymous | NRAS:uc009wgu.3:exon2:c.G37C:p.G13R | 0.312 |
| **974025** | *PHF6* | stopgain | PHF6:uc010nrr.3:exon5:c.C385T:p.R129X | 0.395 |
| **974025** | *RUNX1* | stopgain | RUNX1:uc002yur.1:exon2:c.318_319ins5773:p.P107_P108delinsX | 0.365 |
| **1446307** | *GATA2* | nonsynonymous | GATA2:uc003ekm.3:exon5:c.C911T:p.P304L | 0.155319 |
| **1446307** | *IDH1* | nonsynonymous | IDH1:uc002vcu.3:exon4:c.C394G:p.R132G | 0.519231 |
| **1446307** | *SRSF2* | nonframeshift deletion | SRSF2:uc010wtg.2:exon1:c.279_326del:p.93_109del | 0.32352941 |
| **1794247** | *ASXL1* | frameshift insertion | ASXL1:uc021wbw.1:exon13:c.1926_1927insG:p.G642fs | 0.20134228 |
| **1794247** | *IDH2* | nonsynonymous | IDH2:uc002box.3:exon4:c.G515A:p.R172K | 0.421829 |
| **1799723** | *ASXL1* | stopgain | ASXL1:uc021wbw.1:exon13:c.A2473T:p.K825X | 0.134058 |
| **1799723** | *IDH2* | nonsynonymous | IDH2:uc002box.3:exon4:c.G419A:p.R140Q | 0.264501 |
| **1799723** | *SF3B1* | nonframeshift deletion | SF3B1:uc002uue.3:exon14:c.1854_1856del:p.618_619del | 0.25384615 |
| **1825001** | *IDH2* | nonsynonymous | IDH2:uc002box.3:exon4:c.G419A:p.R140Q | 0.096939 |
| **1825001** | *WT1* | nonsynonymous | WT1:uc001mto.2:exon9:c.G1385C:p.R462P | 0.047817 |
| **1850651** | *IDH2* | nonsynonymous | IDH2:uc002box.3:exon4:c.G419A:p.R140Q | 0.447661 |
| **1850651** | *SRSF2* | nonframeshift deletion | SRSF2:uc010wtg.2:exon1:c.284_307del:p.95_103del | 0.475 |
| **1873085** | *ASXL1* | frameshift deletion | ASXL1:uc021wbw.1:exon13:c.2422delC:p.P808fs | 0.31557377 |
| **1873085** | *DNMT3A* | nonsynonymous | DNMT3A:uc002rgd.4:exon18:c.C2125T:p.P709S | 0.401639 |
| **1873085** | *DNMT3A* | nonsynonymous | DNMT3A:uc002rgd.4:exon21:c.G2409T:p.R803S | 0.395745 |
| **1873085** | *IDH2* | nonsynonymous | IDH2:uc002box.3:exon4:c.G515A:p.R172K | 0.38874 |
| **1873085** | *RUNX1* | splicing | RUNX1:uc010gmw.1:exon6:c.508+2T>A | 0.375 |
| **1956615** | *ASXL1* | frameshift insertion | ASXL1:uc021wbw.1:exon13:c.1926_1927insG:p.G642fs | 0.296875 |
| **1956615** | *IDH2* | nonsynonymous | IDH2:uc002box.3:exon4:c.G419A:p.R140Q | 0.488889 |
| **1956615** | *NRAS* | nonsynonymous | NRAS:uc009wgu.3:exon2:c.G37C:p.G13R | 0.154255 |
| **1956615** | *SRSF2* | nonsynonymous | SRSF2:uc010wtg.2:exon1:c.C284A:p.P95H | 0.470032 |
| **2010157** | *IDH2* | nonsynonymous | IDH2:uc002box.3:exon4:c.A514T:p.R172W | 0.28169 |
| **2150901** | *IDH1* | nonsynonymous | IDH1:uc002vcu.3:exon4:c.G395A:p.R132H | 0.318267 |
| **2150901** | *MGA* | nonsynonymous | MGA:uc010ucy.2:exon17:c.G6125C:p.C2042S | 0.027314 |
| **2150901** | *MGA* | nonsynonymous | MGA:uc010ucy.2:exon17:c.G6202C:p.E2068Q | 0.026354 |
| **2150901** | *MGA* | nonsynonymous | MGA:uc010ucy.2:exon17:c.G6472C:p.A2158P | 0.035541 |
| **2150901** | *RUNX1* | nonsynonymous | RUNX1:uc010gmw.1:exon5:c.G497A:p.R166Q | 0.282534 |
| **2150901** | *RUNX1* | nonframeshift insertion | RUNX1:uc010gmw.1:exon7:c.677_678insGAG:p.S226delinsRS | 0.19285714 |

| | | | | |
|---|---|---|---|---|
| **2150901** | *SRSF2* | nonsynonymous | SRSF2:uc010wtg.2:exon1:c.C284G:p.P95R | 0.31769 |
| **2151503** | *IDH2* | nonsynonymous | IDH2:uc002box.3:exon4:c.G419A:p.R140Q | 0.316872 |
| **2151503** | *MYB* | frameshift insertion | MYB:uc010kgi.3:exon9:c.1109_1110insGATTGGG TGGG:p.S370fs | 0.39597315 |
| **2151503** | *NRAS* | nonsynonymous | NRAS:uc009wgu.3:exon2:c.G34T:p.G12C | 0.017906 |
| **2151503** | *PTPN11* | nonsynonymous | PTPN11:uc001ttx.3:exon3:c.C215A:p.A72D | 0.152174 |
| **2151503** | *RUNX1* | nonsynonymous | RUNX1:uc010gmw.1:exon4:c.C337G:p.P113A | 0.250804 |
| **2151503** | *RUNX1* | nonsynonymous | RUNX1:uc010gmw.1:exon6:c.G592A:p.D198N | 0.230769 |
| **2151503** | *SRSF2* | nonsynonymous | SRSF2:uc010wtg.2:exon1:c.C284A:p.P95H | 0.360324 |
| **2151503** | *TET2* | frameshift deletion | TET2:uc021xqk.1:exon3:c.2747delA:p.Q916fs | 0.04651163 |
| **2152641** | *ASXL1* | stopgain | ASXL1:uc021wbw.1:exon13:c.C2616A:p.C872X | 0.245098 |
| **2152641** | *CEBPA* | frameshift insertion | CEBPA:uc002nun.3:exon1:c.749dupG:p.G250fs | 0.05084746 |
| **2152641** | *IDH2* | nonsynonymous | IDH2:uc002box.3:exon4:c.G419A:p.R140Q | 0.275 |
| **2152641** | *SRSF2* | nonsynonymous | SRSF2:uc010wtg.2:exon1:c.C284T:p.P95L | 0.313725 |
| **2152641** | *STAG2* | nonframeshift deletion | STAG2:uc004eud.3:exon11:c.1010_1012del:p.337_ 338del | 0.49700599 |
| **2163337** | *ASXL1* | stopgain | ASXL1:uc021wbw.1:exon13:c.1772dupA:p.Y591_Q 592delinsX | 0.35048232 |
| **2163337** | *BCOR* | frameshift insertion | BCOR:uc004dep.4:exon14:c.4834dupC:p.L1612fs | 0.07317073 |
| **2163337** | *DNMT3A* | nonsynonymous | DNMT3A:uc002rgd.4:exon6:c.G494A:p.G165D | 0.489914 |
| **2163337** | *IDH2* | nonsynonymous | IDH2:uc002box.3:exon4:c.G419A:p.R140Q | 0.415403 |
| **2163337** | *SRSF2* | nonsynonymous | SRSF2:uc010wtg.2:exon1:c.C284A:p.P95H | 0.480211 |
| **2163337** | *STAG2* | frameshift deletion | STAG2:uc004eud.3:exon5:c.257_261del:p.E86fs | 1 |
| **2210551** | *CEBPA* | stopgain | CEBPA:uc002nun.3:exon1:c.C484T:p.Q162X | 0.526316 |
| **2210551** | *IDH2* | nonsynonymous | IDH2:uc002box.3:exon4:c.G419A:p.R140Q | 0.479833 |
| **2210551** | *MPL* | stopgain | MPL:uc009vwr.3:exon12:c.C1753T:p.R585X | 0.499366 |
| **2210551** | *SRSF2* | nonsynonymous | SRSF2:uc010wtg.2:exon1:c.C284A:p.P95H | 0.487342 |
| **2210551** | *TP53* | nonsynonymous | TP53:uc010cni.2:exon7:c.A704G:p.N235S | 0.43609 |
| **2210551** | *WT1* | stopgain | WT1:uc001mto.2:exon7:c.C1105T:p.R369X | 0.533911 |
| **2219711** | *ASXL1* | frameshift deletion | ASXL1:uc021wbw.1:exon13:c.3633_3636del:p.D12 11fs | 0.38586957 |
| **2219711** | *DNMT3A* | frameshift deletion | DNMT3A:uc002rgd.4:exon18:c.2083_2096del:p.I69 5fs | 0.41666667 |
| **2219711** | *IDH2* | nonsynonymous | IDH2:uc002box.3:exon4:c.G419A:p.R140Q | 0.287335 |
| **2219711** | *RUNX1* | nonsynonymous | RUNX1:uc010gmw.1:exon6:c.G592T:p.D198Y | 0.558304 |
| **2219711** | *SF3B1* | nonsynonymous | SF3B1:uc002uue.3:exon15:c.A2098G:p.K700E | 0.308609 |
| **2219711** | *TET2* | frameshift deletion | TET2:uc021xqk.1:exon3:c.408_429del:p.S136fs | 0.5 |
| **2219711** | *TET2* | frameshift insertion | TET2:uc021xqk.1:exon3:c.623_624insTAATGGTG CTACAGTTTCTGCCTCT:p.P208fs | 0.31284916 |
| **2221709** | *ASXL1* | stopgain | ASXL1:uc021wbw.1:exon13:c.G2113T:p.E705X | 0.263 |
| **2221709** | *DNMT3A* | nonsynonymous | DNMT3A:uc002rgd.3:exon16:c.G1904A:p.R635Q | 0.532 |
| **2221709** | *IDH1* | nonsynonymous | IDH1:uc002vcu.3:exon4:c.C394T:p.R132C | 0.272 |
| **2221709** | *RB1* | stopgain | RB1:uc001vcb.3:exon12:c.C1190G:p.S397X | 0.228 |
| **2221709** | *SMC1A* | splicing | SMC1A:uc011moe.2:exon1:c.109+450G>T | 0.238 |
| **2221709** | *STAG2* | stopgain | STAG2:uc004eud.3:exon8:c.C629A:p.S210X | 0.124 |
| **2222935** | *CEBPA* | nonsynonymous | CEBPA:uc002nun.3:exon1:c.T992A:p.L331Q | 0.274194 |

| | | | | |
|---|---|---|---|---|
| **2222935** | *DNMT3A* | nonsynonymous | DNMT3A:uc002rgd.3:exon10:c.T1232C:p.L411P | 0.253012 |
| **2222935** | *DNMT3A* | nonsynonymous | DNMT3A:uc002rgd.3:exon20:c.A2366C:p.H789P | 0.307317 |
| **2222935** | *IDH2* | nonsynonymous | IDH2:uc002box.3:exon4:c.G419A:p.R140Q | 0.305322 |
| **2222935** | *STAG2* | frameshift deletion | STAG2:uc004eud.3:exon10:c.885delT:p.H295fs | 0.28365385 |
| **2222935** | *ZNF639* | stopgain | ZNF639:uc003fjr.1:exon7:c.G869A:p.W290X | 0.318284 |
| **2244043** | *ASXL1* | frameshift deletion | ASXL1:uc021wbw.1:exon13:c.1888_1910del:p.630_637del | 0.13402062 |
| **2244043** | *IDH1* | nonsynonymous | IDH1:uc002vcu.3:exon4:c.C394T:p.R132C | 0.133333 |
| **2246061** | *ASXL1* | frameshift insertion | ASXL1:uc021wbw.1:exon13:c.1926_1927insG:p.G642fs | 0.15736041 |
| **2246061** | *IDH2* | nonsynonymous | IDH2:uc002box.3:exon4:c.G419A:p.R140Q | 0.053981 |
| **2246061** | *SRSF2* | nonsynonymous | SRSF2:uc010wtg.2:exon1:c.C284A:p.P95H | 0.083624 |
| **2246061** | *STAG2* | frameshift deletion | STAG2:uc004eud.3:exon30:c.3101delG:p.R1034fs | 0.15591398 |
| **2246061** | *TBXA2R* | nonsynonymous | TBXA2R:uc021umv.1:exon2:c.C272G:p.A91G | 0.141026 |
| **2246061** | *TBXA2R* | nonsynonymous | TBXA2R:uc021umv.1:exon2:c.G284C:p.W95S | 0.139764 |
| **2246709** | *IDH2* | nonsynonymous | IDH2:uc002box.3:exon4:c.G515A:p.R172K | 0.41256 |
| **2246709** | *PTPN11* | nonsynonymous | PTPN11:uc001ttx.3:exon3:c.C215T:p.A72V | 0.109091 |
| **2246709** | *RUNX1* | nonsynonymous | RUNX1:uc010gmw.1:exon5:c.G485A:p.R162K | 0.894403 |
| **2246709** | *SF3B1* | nonsynonymous | SF3B1:uc002uue.3:exon15:c.A2098G:p.K700E | 0.444562 |
| **2247707** | *CBL* | nonsynonymous | CBL:uc001pwe.3:exon8:c.C1192T:p.H398Y | 0.094937 |
| **2247707** | *CBL* | nonsynonymous | CBL:uc001pwe.3:exon9:c.G1259T:p.R420L | 0.161702 |
| **2247707** | *DNMT3A* | nonsynonymous | DNMT3A:uc002rgd.3:exon23:c.G2645A:p.R882H | 0.441558 |
| **2247707** | *IDH1* | nonsynonymous | IDH1:uc002vcu.3:exon4:c.G395T:p.R132L | 0.272727 |
| **2247707** | *NPM1* | frameshift insertion | NPM1:uc003mbi.3:exon11:c.859_860insTCTG:p.L287fs | 0.11666667 |
| **2247707** | *WT1* | frameshift deletion | WT1:uc001mto.2:exon8:c.1328_1332del:p.443_444del | 0.06451613 |
| **2247707** | *WT1* | frameshift insertion | WT1:uc001mto.2:exon9:c.1400_1401insGTCAT:p.K467fs | 0.07407407 |
| **2248361** | *ASXL1* | stopgain | ASXL1:uc021wbw.1:exon13:c.C2077T:p.R693X | 0.229 |
| **2248361** | *BCORL1* | frameshift insertion | BCORL1:uc022cdu.1:exon3:c.809_810insAGTTATGT:p.S270fs | 0.248 |
| **2248361** | *DNMT3A* | nonsynonymous | DNMT3A:uc002rgd.3:exon19:c.A2281G:p.M761V | 0.257 |
| **2248361** | *DNMT3A* | nonsynonymous | DNMT3A:uc002rgd.3:exon19:c.T2194C:p.F732L | 0.228 |
| **2248361** | *FLT3* | nonframeshift insertion | FLT3:uc010tdn.2:exon14:c.1818_1819insGGGGAGGGGTGAATATGATCTCAAATGGGAGTTTCC:p.P606delinsPGRGEYDLKWEFP | 0.13 |
| **2248361** | *IDH1* | nonsynonymous | IDH1:uc002vcu.3:exon4:c.C394T:p.R132C | 0.237 |
| **2248361** | *RUNX1* | frameshift deletion | RUNX1:uc010gmv.3:exon9:c.977delA:p.D326fs | 0.204 |
| **2248361** | *RUNX1* | frameshift insertion | RUNX1:uc010gmw.1:exon5:c.352_353insTTTAG:p.V118fs | 0.228 |
| **2260553** | *DNMT3A* | nonsynonymous | DNMT3A:uc002rgd.3:exon23:c.G2645A:p.R882H | 0.391403 |
| **2260553** | *FLT3* | nonframeshift insertion | FLT3:uc010tdn.2:exon14:c.1780_1781insATGAAAGCCAGCTACAGATGGTACAGGTGACCGGCTCCTCAGATAATGAGTACTTCTACGTTGATT:p.F594delinsYESQLQMVQVTGSSDNEYFYVDF | 0.38797814 |
| **2260553** | *IDH2* | nonsynonymous | IDH2:uc002box.3:exon4:c.G419A:p.R140Q | 0.367241 |
| **2260553** | *NPM1* | frameshift insertion | NPM1:uc003mbi.3:exon11:c.859_860insTCTG:p.L287fs | 0.3627451 |
| **2260671** | *IDH2* | nonsynonymous | IDH2:uc002box.3:exon4:c.G419A:p.R140Q | 0.21 |
| **2260671** | *NPM1* | frameshift | NPM1:uc003mbi.3:exon11:c.859_860insTCTG:p.L287fs | 0.09 |
| **2260671** | *SRSF2* | nonsynonymous | SRSF2:uc010wtg.2:exon1:c.C284T:p.P95L | 0.221 |

| | | | | |
|---|---|---|---|---|
| **2263055** | *IDH1* | nonsynonymous | IDH1:uc002vcu.3:exon4:c.C394T:p.R132C | 0.061657 |
| **2263055** | *NRAS* | nonsynonymous | NRAS:uc009wgu.3:exon2:c.G35C:p.G12A | 0.019116 |
| **2263055** | *TP53* | nonsynonymous | TP53:uc010cni.2:exon5:c.C380T:p.S127F | 0.983607 |
| **2264225** | *DNMT3A* | splicing/nonsynonymous | DNMT3A:uc002rgd.3:exon21:c.A2477G:p.K826R | 0.87 |
| **2264225** | *IDH1* | nonsynonymous | IDH1:uc002vcu.3:exon4:c.C394T:p.R132C | 0.333333 |
| **2264225** | *SETD2* | splicing/nonsynonymous | SETD2:uc003cqv.3:exon21:c.T7634A:p.M2545K | 0.402878 |
| **2264225** | *SETD2* | nonsynonymous | SETD2:uc003cqv.3:exon21:c.T7636A:p.S2546T | 0.395683 |
| **2264225** | *SETD2* | frameshift deletion | SETD2:uc003cqv.3:exon3:c.574_575del:p.192_192del | 0.26701571 |
| **2264727** | *DNMT3A* | nonsynonymous | DNMT3A:uc002rgd.4:exon22:c.A2512G:p.N838D | 0.348941 |
| **2264727** | *IDH1* | nonsynonymous | IDH1:uc002vcu.3:exon4:c.C394T:p.R132C | 0.129 |
| **2264727** | *MGA* | nonsynonymous | MGA:uc010ucy.2:exon24:c.T8138G:p.L2713W | 0.509989 |
| **2264727** | *NPM1* | frameshift insertion | NPM1:uc003mbi.3:exon11:c.859_860insTCTG:p.L287fs | 0.24528302 |
| **2264727** | *NRAS* | nonsynonymous | NRAS:uc009wgu.3:exon2:c.G38A:p.G13D | 0.310308 |
| **2264727** | *RAD21* | frameshift deletion | RAD21:uc003yod.3:exon3:c.219_228del:p.Y73fs | 0.51351351 |
| **2264727** | *TP53* | nonsynonymous | TP53:uc010cni.2:exon7:c.A704G:p.N235S | 0.480769 |
| **2280935** | *ASXL1* | frameshift deletion | ASXL1:uc021wbw.1:exon13:c.2057_2058del:p.K686fs | 0.18617021 |
| **2280935** | *IDH2* | nonsynonymous | IDH2:uc002box.3:exon4:c.G419A:p.R140Q | 0.126415 |
| **2281469** | *ASXL1* | frameshift deletion | ASXL1:uc021wbw.1:exon13:c.1888_1910del:p.630_637del | 0.32380952 |
| **2281469** | *BCOR* | stopgain | BCOR:uc004dep.4:exon8:c.C3613T:p.Q1205X | 0.021053 |
| **2281469** | *DNMT3A* | nonsynonymous | DNMT3A:uc002rgd.3:exon23:c.C2644T:p.R882C | 0.536913 |
| **2281469** | *IDH1* | nonsynonymous | IDH1:uc002vcu.3:exon4:c.C394T:p.R132C | 0.419355 |
| **2281469** | *RUNX1* | nonsynonymous | RUNX1:uc010gmw.1:exon5:c.G497A:p.R166Q | 0.46114 |
| **2281469** | *SRSF2* | nonsynonymous | SRSF2:uc010wtg.2:exon1:c.C284A:p.P95H | 0.492991 |
| **2283265** | *IDH2* | nonsynonymous | IDH2:uc002box.3:exon4:c.G419A:p.R140Q | 0.401544 |
| **2292873** | *DNMT3A* | frameshift deletion | DNMT3A:uc002rgd.3:exon17:c.1979_1988del:p.660_663del | 0.57291667 |
| **2292873** | *DNMT3A* | stopgain | DNMT3A:uc002rgd.3:exon8:c.G980A:p.W327X | 0.448485 |
| **2292873** | *IDH1* | nonsynonymous | IDH1:uc002vcu.3:exon4:c.C394T:p.R132C | 0.446115 |
| **2292915** | *ASXL1* | frameshift deletion | ASXL1:uc021wbw.1:exon13:c.1773_1774del:p.Y591fs | 0.33090909 |
| **2292915** | *IDH2* | nonsynonymous | IDH2:uc002box.3:exon4:c.G419A:p.R140Q | 0.412884 |
| **2292915** | *U2AF1* | nonsynonymous | U2AF1:uc002zcz.1:exon7:c.A251C:p.Q84P | 0.459119 |
| **2295595** | *DNMT3A* | nonsynonymous | DNMT3A:uc002rgd.3:exon23:c.G2645A:p.R882H | 0.381982 |
| **2295595** | *IDH2* | nonsynonymous | IDH2:uc002box.3:exon4:c.G419A:p.R140Q | 0.328025 |
| **2295595** | *PHF6* | frameshift deletion | PHF6:uc010nrr.3:exon7:c.676delG:p.G226fs | 0.1509434 |
| **2295595** | *SMC1A* | splicing/nonsynonymous | SMC1A:uc011moe.2:exon16:c.G2354A:p.R785H | 0.695896 |
| **2295595** | *SRSF2* | nonsynonymous | SRSF2:uc010wtg.2:exon1:c.C284A:p.P95H | 0.389655 |
| **2296723** | *ASXL1* | frameshift insertion | ASXL1:uc021wbw.1:exon13:c.1926_1927insG:p.G642fs | 0.20714286 |
| **2296723** | *DNMT3A* | splicing | DNMT3A:uc002rgd.3:exon13:c.1474+1G>T | 0.068182 |
| **2296723** | *DNMT3A* | nonsynonymous | DNMT3A:uc002rgd.3:exon23:c.G2645A:p.R882H | 0.408333 |
| **2296723** | *IDH2* | nonsynonymous | IDH2:uc002box.3:exon4:c.G515A:p.R172K | 0.452532 |
| **2297625** | *ASXL1* | frameshift insertion | ASXL1:uc021wbw.1:exon13:c.1926_1927insG:p.G642fs | 0.0990991 |

| | | | | |
|---|---|---|---|---|
| **2297625** | *IDH2* | nonsynonymous | IDH2:uc002box.3:exon4:c.G419A:p.R140Q | 0.083045 |
| **2297625** | *PHF6* | frameshift deletion | PHF6:uc010nrr.3:exon3:c.221delT:p.I74fs | 0.17948718 |
| **2297625** | *RUNX1* | splicing | RUNX1:uc010gmw.1:exon6:c.508+2T>C | 0.289655 |
| **2297625** | *SRSF2* | nonsynonymous | SRSF2:uc010wtg.2:exon1:c.C284A:p.P95H | 0.300283 |
| **2297625** | *STAG2* | frameshift insertion | STAG2:uc004eud.3:exon28:c.2797_2798insA:p.E933fs | 0.06818182 |
| **2297625** | *TET2* | nonsynonymous | TET2:uc011cez.2:exon9:c.G4145C:p.G1382A | 0.055901 |
| **2297625** | *TET2* | frameshift deletion | TET2:uc021xqk.1:exon3:c.1222delC:p.P408fs | 0.27319588 |
| **2297707** | *BCOR* | stopgain | BCOR:uc004deq.4:exon4:c.C1971G:p.Y657X | 0.1 |
| **2297707** | *DNMT3A* | nonframeshift insertion | DNMT3A:uc002rgd.3:exon17:c.1983_1984insGGGCATTCAGGTGGACCGCTACAT:p.I661delinsMGIQVDRYI | 0.294 |
| **2297707** | *DNMT3A* | nonsynonymous | DNMT3A:uc002rgd.3:exon23:c.G2645A:p.R882H | 0.07 |
| **2297707** | *IDH1* | nonsynonymous | IDH1:uc002vcu.3:exon4:c.C394T:p.R132C | 0.066 |
| **2297941** | *DNMT3A* | nonsynonymous | DNMT3A:uc002rgd.3:exon23:c.G2645A:p.R882H | 0.349451 |
| **2297941** | *IDH1* | nonsynonymous | IDH1:uc002vcu.3:exon4:c.G395T:p.R132L | 0.361111 |
| **2307799** | *DNMT3A* | nonsynonymous | DNMT3A:uc002rgd.3:exon16:c.C1915T:p.L639F | 0.05 |
| **2307799** | *DNMT3A* | nonsynonymous | DNMT3A:uc002rgd.3:exon23:c.C2611G:p.P871A | 0.34 |
| **2307799** | *IDH2* | nonsynonymous | IDH2:uc002box.3:exon4:c.G515A:p.R172K | 0.32 |
| **2337831** | *BCORL1* | stopgain | BCORL1:uc022cdu.1:exon3:c.2406_2407insTAGGGGA:p.L802_S803delinsLX | 0.20526316 |
| **2337831** | *IDH2* | nonsynonymous | IDH2:uc002box.3:exon4:c.G419A:p.R140Q | 0.447531 |
| **2337831** | *RUNX1* | frameshift insertion | RUNX1:uc010gmv.3:exon9:c.1252_1253insC:p.M418fs | 0.27131783 |
| **2337831** | *SRSF2* | nonsynonymous | SRSF2:uc010wtg.2:exon1:c.C284T:p.P95L | 0.40201 |
| **2361361** | *IDH2* | nonsynonymous | IDH2:uc002box.3:exon4:c.G515A:p.R172K | 0.017354 |
| **2361361** | *MED12* | nonsynonymous | MED12:uc011mpq.1:exon23:c.A3332G:p.N1111S | 0.013447 |
| **2361405** | *ASXL1* | frameshift insertion | ASXL1:uc021wbw.1:exon13:c.1926_1927insG:p.G642fs | 0.15384615 |
| **2361405** | *IDH2* | nonsynonymous | IDH2:uc002box.3:exon4:c.G419A:p.R140Q | 0.348135 |
| **2361405** | *SRSF2* | nonsynonymous | SRSF2:uc010wtg.2:exon1:c.C284A:p.P95H | 0.336879 |
| **2361767** | *DNMT3A* | nonsynonymous | DNMT3A:uc002rgd.3:exon23:c.G2645A:p.R882H | 0.245161 |
| **2361767** | *IDH1* | nonsynonymous | IDH1:uc002vcu.3:exon4:c.C394T:p.R132C | 0.061 |
| **2361767** | *MYB* | stopgain | MYB:uc003qfq.3:exon10:c.C1553A:p.S518X | 0.055556 |
| **2366711** | *DNMT3A* | nonsynonymous | DNMT3A:uc002rgd.3:exon23:c.G2645A:p.R882H | 0.155702 |
| **2366711** | *IDH2* | nonsynonymous | IDH2:uc002box.3:exon4:c.G515A:p.R172K | 0.138298 |
| **2366711** | *PHF6* | frameshift insertion | PHF6:uc011mvk.2:exon2:c.93dupA:p.L31fs | 0.11267606 |
| **2366711** | *RUNX1* | frameshift insertion | RUNX1:uc010gmw.1:exon6:c.548dupC:p.P183fs | 0.17991632 |
| **2366711** | *XPO1* | nonsynonymous | XPO1:uc010ypn.2:exon19:c.C2170T:p.L724F | 0.04401 |
| **2370233** | *ASXL1* | frameshift deletion | ASXL1:uc021wbw.1:exon13:c.2032_2033del:p.R678fs | 0.14218009 |
| **2370233** | *CEBPA* | frameshift deletion | CEBPA:uc002nun.3:exon1:c.273delC:p.A91fs | 0.06410256 |
| **2370233** | *DNMT3A* | nonsynonymous | DNMT3A:uc002rgd.4:exon14:c.G1628C:p.G543A | 0.12967 |
| **2370233** | *IDH1* | nonsynonymous | IDH1:uc002vcu.3:exon4:c.C394T:p.R132C | 0.135011 |
| **2370233** | *NPM1* | frameshift insertion | NPM1:uc003mbi.3:exon11:c.859_860insTCTG:p.L287fs | 0.11827957 |
| **2370233** | *SETD2* | stopgain | SETD2:uc003cqv.3:exon3:c.C3997T:p.Q1333X | 0.059805 |
| **2370759** | *BCOR* | stopgain | BCOR:uc004dep.4:exon10:c.G4288T:p.E1430X | 0.235294 |

| | | | | |
|---|---|---|---|---|
| **2370759** | *CBL* | nonframeshift deletion | CBL:uc001pwe.3:exon8:c.1096_1227del:p.366_409del | 0.06862745 |
| **2370759** | *DNMT3A* | nonsynonymous | DNMT3A:uc002rgd.3:exon23:c.G2645A:p.R882H | 0.281538 |
| **2370759** | *IDH2* | nonsynonymous | IDH2:uc002box.3:exon4:c.G419A:p.R140Q | 0.215926 |
| **2370759** | *RUNX1* | frameshift deletion | RUNX1:uc010gmv.3:exon9:c.1096_1103del:p.366_368del | 0.22222222 |
| **2370759** | *RUNX1* | stopgain | RUNX1:uc010gmv.3:exon9:c.C1209A:p.Y403X | 0.021786 |
| **2370759** | *SRSF2* | nonsynonymous | SRSF2:uc010wtg.2:exon1:c.C284A:p.P95H | 0.259155 |
| **2370759** | *TET2* | frameshift deletion | TET2:uc021xqk.1:exon3:c.2936delG:p.R979fs | 0.06325301 |
| **2370759** | *TET2* | stopgain | TET2:uc021xqk.1:exon3:c.G655T:p.E219X | 0.325658 |
| **2382431** | *IDH2* | nonsynonymous | IDH2:uc002box.3:exon4:c.G419A:p.R140Q | 0.367857 |
| **2382431** | *SMC1A* | stopgain | SMC1A:uc011moe.2:exon18:c.G2497T:p.E833X | 0.776515 |
| **2383235** | *DNMT3A* | frameshift deletion | DNMT3A:uc002rgd.3:exon21:c.2426_2427del:p.809_809del | 0.39189189 |
| **2383235** | *IDH2* | nonsynonymous | IDH2:uc002box.3:exon4:c.G419A:p.R140Q | 0.19244 |
| **2394529** | *ASXL1* | stopgain | ASXL1:uc021wbw.1:exon13:c.C2309G:p.S770X | 0.427918 |
| **2394529** | *DNMT3A* | frameshift deletion | DNMT3A:uc002rgd.3:exon11:c.1372delC:p.R458fs | 0.19125683 |
| **2394529** | *DNMT3A* | frameshift deletion | DNMT3A:uc002rgd.3:exon19:c.2194_2207del:p.732_736del | 0.26829268 |
| **2394529** | *IDH1* | nonsynonymous | IDH1:uc002vcu.3:exon4:c.C394T:p.R132C | 0.304348 |
| **2394529** | *IDH2* | nonsynonymous | IDH2:uc002box.3:exon4:c.G419A:p.R140Q | 0.194203 |
| **2394529** | *U2AF1* | nonsynonymous | U2AF1:uc002zcz.1:exon7:c.A251G:p.Q84R | 0.239286 |
| **2410163** | *IDH1* | nonsynonymous | IDH1:uc002vcu.3:exon4:c.C394T:p.R132C | 0.372685 |
| **2412147** | *DNMT3A* | nonsynonymous | DNMT3A:uc002rgd.3:exon18:c.G2096A:p.G699D | 0.554 |
| **2412147** | *IDH2* | nonsynonymous | IDH2:uc002box.3:exon4:c.G515A:p.R172K | 0.322 |
| **2419959** | *ASXL1* | frameshift deletion | ASXL1:uc021wbw.1:exon13:c.3633_3636del:p.D1211fs | 0.07303371 |
| **2419959** | *IDH2* | nonsynonymous | IDH2:uc002box.3:exon4:c.G515A:p.R172K | 0.050725 |
| **2426545** | *ASXL1* | frameshift deletion | ASXL1:uc021wbw.1:exon13:c.2818delT:p.L940fs | 0.26133333 |
| **2426545** | *IDH1* | nonsynonymous | IDH1:uc002vcu.3:exon4:c.C394T:p.R132C | 0.096026 |
| **2426545** | *NF1* | frameshift insertion | NF1:uc002hgh.3:exon5:c.504_505insCCCATCAC:p.S168fs | 0.1589404 |
| **2426545** | *RUNX1* | frameshift insertion | RUNX1:uc010gmv.3:exon9:c.1028_1029insTGATC:p.S343fs | 0.39772727 |
| **2426545** | *RUNX1* | frameshift insertion | RUNX1:uc010gmw.1:exon6:c.540dupC:p.T181fs | 0.2688172 |
| **2426545** | *SRSF2* | nonframeshift insertion | SRSF2:uc010wtg.2:exon1:c.283_284insGCG:p.P95delinsRA | 0.17021277 |
| **2430935** | *IDH1* | nonsynonymous | IDH1:uc002vcu.3:exon4:c.C394T:p.R132C | 0.221154 |
| **2430935** | *SRSF2* | nonsynonymous | SRSF2:uc010wtg.2:exon1:c.C284A:p.P95H | 0.187234 |
| **2432389** | *ASXL1* | frameshift insertion | ASXL1:uc021wbw.1:exon13:c.1926_1927insG:p.G642fs | 0.09240924 |
| **2432389** | *DNMT3A* | frameshift insertion | DNMT3A:uc002rgd.3:exon19:c.2303_2304insCGTTAGTGACAAGAGGG:p.D768fs | 0.24669604 |
| **2432389** | *IDH1* | nonsynonymous | IDH1:uc002vcu.3:exon4:c.C394T:p.R132C | 0.223192 |
| **2432389** | *PHF6* | stopgain | PHF6:uc010nrr.3:exon8:c.G811T:p.E271X | 0.069277 |
| **2432389** | *U2AF1* | nonsynonymous | U2AF1:uc002zcz.1:exon7:c.A251C:p.Q84P | 0.339192 |
| **2450523** | *ASXL1* | frameshift insertion | ASXL1:uc021wbw.1:exon13:c.1926_1927insG:p.G642fs | 0.27699531 |
| **2450523** | *CEBPA* | frameshift insertion | CEBPA:uc002nun.3:exon1:c.68dupC:p.P23fs | 0.04166667 |
| **2450523** | *CEBPA* | frameshift insertion | CEBPA:uc002nun.3:exon1:c.81_82insCC:p.S28fs | 0.07692308 |
| **2450523** | *CEBPA* | stopgain | CEBPA:uc002nun.3:exon1:c.C324G:p.Y108X | 0.294118 |

| | | | | |
|---|---|---|---|---|
| **2450523** | *CEBPA* | nonsynonymous | CEBPA:uc002nun.3:exon1:c.C970G:p.L324V | 0.008347 |
| **2450523** | *GATA2* | frameshift insertion | GATA2:uc003ekm.3:exon3:c.72dupA:p.H25fs | 0.15 |
| **2450523** | *IDH1* | nonsynonymous | IDH1:uc002vcu.3:exon4:c.C394T:p.R132C | 0.184676 |
| **2450523** | *RUNX1* | splicing | RUNX1:uc010gmw.1:exon8:c.806-2A>G | 0.233716 |
| **2450523** | *SRSF2* | nonsynonymous | SRSF2:uc010wtg.2:exon1:c.C284A:p.P95H | 0.440741 |
| **2450523** | *TET2* | frameshift deletion | TET2:uc021xqk.1:exon3:c.1895delA:p.Q632fs | 0.319202 |
| **2463247** | *ASXL1* | frameshift insertion | ASXL1:uc021wbw.1:exon13:c.1926_1927insG:p.G642fs | 0.22807018 |
| **2463247** | *IDH1* | nonsynonymous | IDH1:uc002vcu.3:exon4:c.C394T:p.R132C | 0.26087 |
| **2463247** | *KRAS* | nonsynonymous | KRAS:uc001rgq.1:exon4:c.C437T:p.A146V | 0.021858 |
| **2463247** | *NRAS* | nonsynonymous | NRAS:uc009wgu.3:exon2:c.G34C:p.G12R | 0.016194 |
| **2463247** | *NRAS* | nonsynonymous | NRAS:uc009wgu.3:exon3:c.T190G:p.Y64D | 0.104972 |
| **2463247** | *RUNX1* | nonsynonymous | RUNX1:uc010gmw.1:exon6:c.G602A:p.R201Q | 0.475771 |
| **2463247** | *SRSF2* | nonsynonymous | SRSF2:uc010wtg.2:exon1:c.C284A:p.P95H | 0.462963 |
| **2480485** | *IDH2* | nonsynonymous | IDH2:uc002box.3:exon4:c.G419A:p.R140Q | 0.473214 |
| **2480485** | *JAK2* | nonsynonymous | JAK2:uc003ziw.3:exon14:c.G1849T:p.V617F | 0.997859 |
| **2480485** | *RUNX1* | frameshift insertion | RUNX1:uc010gmv.3:exon9:c.1023dupC:p.I342fs | 0.16071429 |
| **2480485** | *RUNX1* | nonsynonymous | RUNX1:uc010gmw.1:exon6:c.G592A:p.D198N | 0.538018 |
| **2480485** | *SRSF2* | nonsynonymous | SRSF2:uc010wtg.2:exon1:c.C284T:p.P95L | 0.502058 |
| **2498789** | *ASXL1* | frameshift insertion | ASXL1:uc021wbw.1:exon13:c.1926_1927insG:p.G642fs | 0.0621118 |
| **2498789** | *IDH2* | nonsynonymous | IDH2:uc002box.3:exon4:c.G419A:p.R140Q | 0.075251 |
| **2498789** | *SRSF2* | nonsynonymous | SRSF2:uc010wtg.2:exon1:c.C284A:p.P95H | 0.104803 |
| **2502699** | *IDH1* | nonsynonymous | IDH1:uc002vcu.3:exon4:c.C394T:p.R132C | 0.488372 |
| **2502699** | *SRSF2* | nonsynonymous | SRSF2:uc010wtg.2:exon1:c.C284G:p.P95R | 0.400593 |
| **2502699** | *TET2* | stopgain | TET2:uc021xqk.1:exon3:c.C2728T:p.Q910X | 0.006897 |
| **2512641** | *DNMT3A* | frameshift deletion | DNMT3A:uc002rgd.3:exon6:c.503delG:p.G168fs | 0.28742515 |
| **2512641** | *GATA2* | frameshift deletion | GATA2:uc003ekm.3:exon4:c.625delG:p.D209fs | 0.20164609 |
| **2512641** | *IDH1* | nonsynonymous | IDH1:uc002vcu.3:exon4:c.G395A:p.R132H | 0.24826 |
| **2583919** | *DNMT3A* | splicing | DNMT3A:uc002rgd.3:exon11:c.1429+1G>A | 0.284738 |
| **2583919** | *IDH2* | nonsynonymous | IDH2:uc002box.3:exon4:c.G419A:p.R140Q | 0.301923 |
| **2583919** | *MGA* | stopgain | MGA:uc010ucz.2:exon9:c.C3157T:p.R1053X | 0.300842 |
| **2583919** | *NPM1* | frameshift insertion | NPM1:uc003mbi.3:exon11:c.859_860insTCTG:p.L287fs | 0.25396825 |
| **2616045** | *BCOR* | frameshift insertion | BCOR:uc004deq.4:exon4:c.1218dupG:p.P407fs | 0.21969697 |
| **2616045** | *DNMT3A* | nonsynonymous | DNMT3A:uc002rgd.3:exon23:c.G2645A:p.R882H | 0.422785 |
| **2616045** | *IDH2* | nonsynonymous | IDH2:uc002box.3:exon4:c.G419A:p.R140Q | 0.412556 |
| **2616045** | *RUNX1* | nonframeshift deletion | RUNX1:uc010gmw.1:exon4:c.341_343del:p.114_115del | 0.92307692 |
| **2616045** | *SRSF2* | nonsynonymous | SRSF2:uc010wtg.2:exon1:c.C284T:p.P95L | 0.485149 |
| **2620771** | *BCOR* | stopgain | BCOR:uc004dep.4:exon8:c.G3707A:p.W1236X | 0.903509 |
| **2620771** | *DNMT3A* | nonsynonymous | DNMT3A:uc002rgd.3:exon19:c.C2245G:p.R749G | 0.426901 |
| **2620771** | *DNMT3A* | nonsynonymous | DNMT3A:uc002rgd.3:exon23:c.T2726C:p.F909S | 0.445596 |
| **2620771** | *IDH1* | nonsynonymous | IDH1:uc002vcu.3:exon4:c.C394T:p.R132C | 0.285714 |

| | | | | |
|---|---|---|---|---|
| **2620771** | *NF1* | splicing | NF1:uc002hgh.3:exon13:c.1527+3A>T | 0.29906542 |
| **2620771** | *SRSF2* | nonsynonymous | SRSF2:uc010wtg.2:exon1:c.C284T:p.P95L | 0.40583 |
| **2620771** | *STAG2* | frameshift insertion | STAG2:uc004eud.3:exon22:c.2118_2119insGATTT ATTTGCTTGTAATTA:p.W706fs | 1 |
| **4128869** | *BCOR* | stopgain | BCOR:uc004dep.4:exon9:c.C3814T:p.Q1272X | 0.84375 |
| **4128869** | *DNMT3A* | nonsynonymous | DNMT3A:uc002rgd.3:exon23:c.G2645A:p.R882H | 0.413333 |
| **4128869** | *IDH1* | nonsynonymous | IDH1:uc002vcu.3:exon4:c.C394T:p.R132C | 0.408367 |
| **4128869** | *RUNX1* | frameshift deletion | RUNX1:uc010gmw.1:exon7:c.648delC:p.P216fs | 0.28776978 |
| **4128869** | *SRSF2* | nonframeshift insertion | SRSF2:uc010wtg.2:exon1:c.283_284insGCC:p.P95d elinsRP | 0.32022472 |
| **4235533** | *FLT3* | nonsynonymous | FLT3:uc010tdn.2:exon11:c.C1352T:p.S451F | 0.446084 |
| **4235533** | *IDH1* | nonsynonymous | IDH1:uc002vcu.3:exon4:c.G395A:p.R132H | 0.459406 |
| **4235533** | *STAG2* | frameshift deletion | STAG2:uc004eud.3:exon4:c.90delC:p.I30fs | 0.57241379 |
| **4235533** | *TET2* | nonsynonymous | TET2:uc011cez.2:exon7:c.G3877A:p.A1293T | 0.433007 |
| **4235533** | *WT1* | frameshift deletion | WT1:uc001mtq.2:exon2:c.720delC:p.P240fs | 0.31679389 |
| **4259035** | *BCOR* | splicing | BCOR:uc004deq.4:exon10:c.4326+2T>G | 0.124306 |
| **4259035** | *HIST1H2BL* | nonsynonymous | HIST1H2BL:uc003njl.3:exon1:c.C255G:p.N85K | 0.026154 |
| **4259035** | *IDH1* | nonsynonymous | IDH1:uc002vcu.3:exon4:c.C394T:p.R132C | 0.241379 |
| **4259035** | *WT1* | frameshift insertion | WT1:uc001mto.2:exon10:c.1527dupA:p.L510fs | 0.20819113 |

**Supplemental Methods. Targeted deep sequencing data analysis.**

Raw sequencing data from the Illumina platform were converted to a fastq format and aligned to the reference genome (hg19) using the Burroughs-Wheeler Aligner (BWA). BWA is using MEM mode with following parameters: -k 31 -T 100 -t 8 -M. The aligned BAM files were subjected to mark duplication, re-alignment, and re-calibration using Picard and GATK with default parameters (https://www.broadinstitute.org/gatk/guide/best-practices?bpm=DNAseq, last accessed 9/29/2016). Preprocessed BAM files were then analyzed to detect single nucleotide variants (SNV) and small insertions and deletions (indels) using MuTect (https://pubmed.ncbi.nlm.nih.gov/19451168/) and Pindel (https://pubmed.ncbi.nlm.nih.gov/19561018/) algorithms, respectively.
We performed a series of filtering and annotation to identify high-confidence driver mutations. First, variants with low quality sequencing data were filtered out. Specifically, variants matching one or more of the following criteria were considered of low quality and therefore filtered out from further analysis: 1) tumor coverage < 15X, 2) tumor allele frequency < 5%, and 3) normal allele frequency >= 1% and 0% for SNVs and INDELs, respectively. Second, only variants which would introduce an obvious protein-coding change were kept for further analysis. Specifically, variant with an ANNOVAR annotation of non-synonymous, stop-gain, stop-loss, splicing, frameshift insertion, frameshift deletion, nonframeshift insertion or nonframeshift deletion was considered to be able to introduce an obvious protein-coding change and were therefore kept for further analysis. Third, common polymorphisms were removed to reduce the load of possible germline contamination due to the absence of matched normal. Specifically, a series of public variant database including the 1000 Genome Database (http://www.1000genomes.org/), ESP6500 Database (http://evs.gs.washington.edu/EVS/), dbSNP ver.129 (http://www.ncbi.nlm.nih.gov/SNP/), and Exome Aggregation Consortium database (http://exac.broadinstitute.org/), were utilized. Variant with a population frequency of 0.14% or more in any of the databases was considered possible germline polymorphism and was therefore removed from further analysis. Finally, a hierarchical classification system was developed to assign confidence level for each remaining variant in order to facilitate the identification of putative driver mutations. Specifically, each variant was classified based on the following hierarchical order and was assigned a confidence level corresponding to its rank in the system: 1) Confirmed somatic mutation based on COSMIC database (version 81), 2) loss-of-function mutation such as splicing, stop-gain, stop-loss and frameshift mutation in tumor suppressor genes, 3) variant which resides in the same position as a confirmed somatic mutation according to the COSMIC database, 4) recurrent variant which resides within three amino acids away from a confirmed somatic mutation according to the COSMIC database and was predicted to be damaging by in-silico function prediction algorithms. The final annotated variant list was then further analyzed by manual inspection in order to identify putative driver mutation.
